# Origins of 1/f-like tissue oxygenation fluctuations in the murine cortex

**DOI:** 10.1101/2020.09.18.303164

**Authors:** Qingguang Zhang, Kyle W. Gheres, Patrick J. Drew

**Affiliations:** Department of Engineering Science and Mechanics, The Pennsylvania State University, University Park, PA, USA; Graduate Program in Molecular Cellular and Integrative Biosciences, The Pennsylvania State University, University Park, PA, USA; Department of Neurosurgery, The Pennsylvania State University, University Park, PA, USA; Department of Biomedical Engineering, The Pennsylvania State University, University Park, PA, USA

**Keywords:** tissue oxygen, laminar electrophysiology, power spectrum, 1/f

## Abstract

The concentration of oxygen in the brain spontaneously fluctuates, and the power distribution in these fluctuations has 1/f-like dynamics. Though these oscillations have been interpreted as being driven by neural activity, the origins of these 1/f-like oscillations is not well understood. Here, to gain insight of the origin of the 1/f-like oxygen fluctuations, we investigated the dynamics of tissue oxygenation and neural activity in awake behaving mice. We found that oxygen signal recorded from the cortex of mice had 1/f-like spectra. However, band-limited power in the local field potential, did not show corresponding 1/f-like fluctuations. When local neural activity was suppressed, the 1/f-like fluctuations in oxygen concentration persisted. Two-photon measurements of erythrocyte spacing fluctuations (‘stalls’) and mathematical modelling show that stochastic fluctuations in erythrocyte flow and stalling could underlie 1/f-like dynamics in oxygenation. These results show discrete nature of erythrocytes and their irregular flow, rather than neural activity, could drive 1/f-like fluctuations in tissue oxygenation.

## Introduction

Fluctuations in oxygen tension are ubiquitous throughout the body, and are found in muscle tissue and tumors [1], in the retina [2, 3], in the carotid artery [4], and in the cortex [5–12]. Despite their ubiquity, relatively little is understood about the origin of these oxygen fluctuations. While some of these fluctuations are driven by fluctuations in respiration, such as the breathing rate and intensity [4, 13–20], fluctuations in oxygen concentration are present covering a wide range of frequency, not just at the respiration frequency, with most of the power concentrated at lower (< 0.1 Hz) frequencies [1,2, 5, 7–9]. The power spectrum of oxygen concentrations in many tissues show a “1/f-like” behavior, that is, the power at any given frequency *f* is proportional to 1/*f^β^*, where the exponent *β* is usually between 1 and 2 [8]. We refer to these oscillations as being 1/f-like because they are only characterized within a limited frequency region (≥ 0.01 Hz and ≤ 1 Hz). While many biological processes have been shown to exhibit 1/f-like dynamics, a process can only be said to be 1/f if there is data over at least two orders of magnitude in both the abscissa and ordinate [21], a criterion that only a few studies meet [8]. In contrast, white noise has a constant power across frequencies, which when fitted with a power law gives a *β* close to 0 (**Figure 1**a). In both cases, there can be “extra” spectral power concentrated in a single band, leading to a “bump” in the spectrum (**Figure 1**b). Measurements of tissue oxygenation in primates show a clear, statistically robust 1/f-like power spectra, with an additional peak near 0.1 Hz [8].

**Figure 1.**
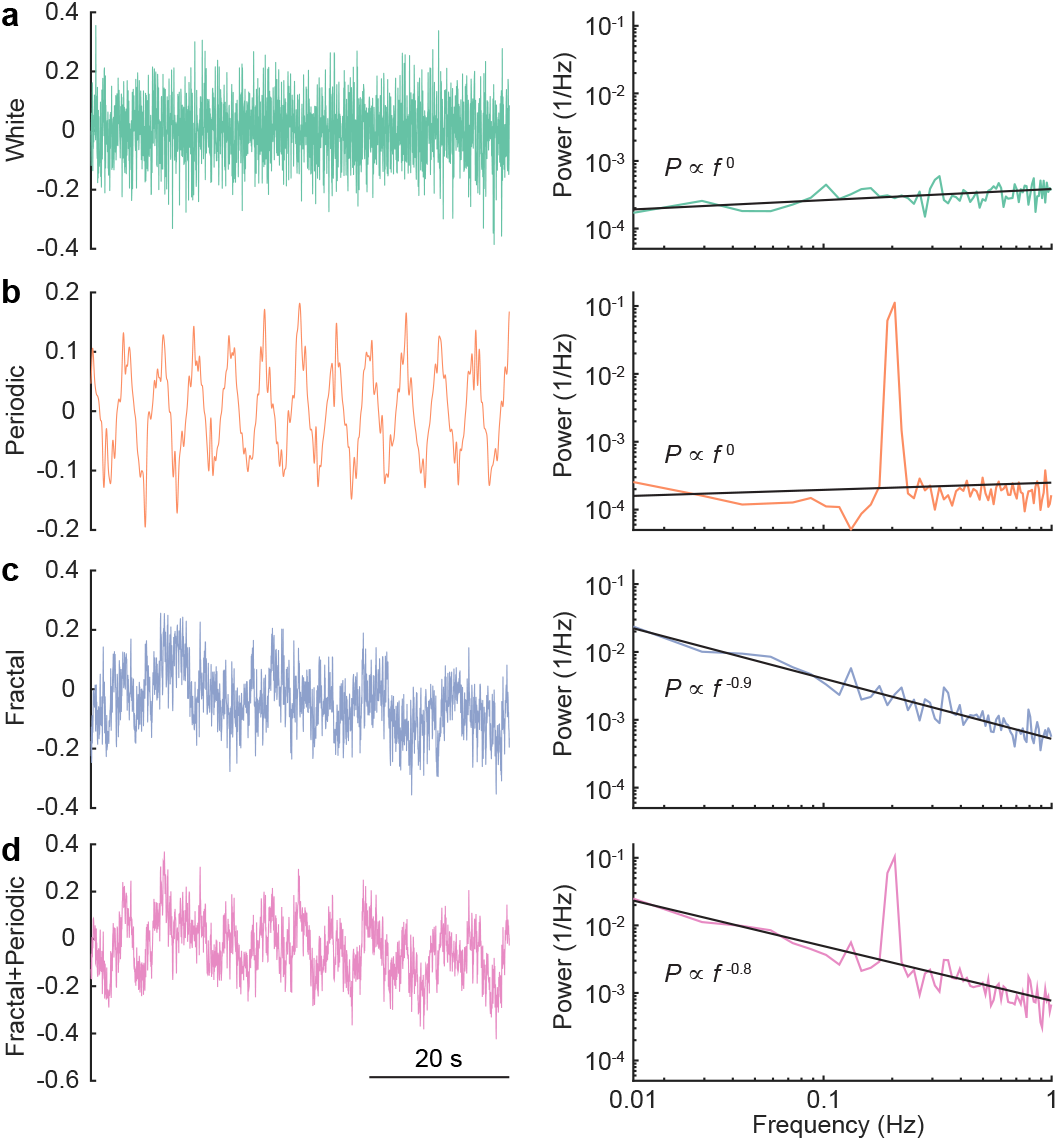
Illustration of white noise, periodic and 1/f-like signal. (**a**) An example of white noise (left) and its power spectrum (right). The solid black line denotes the linear regression fit. (**b**) An example of a periodic signal with peak frequency centered at 0.2 Hz (left) and its power spectrum (right). (**c**) An example of fractional Gaussian noise (i.e., 1/f-like) with Hurst exponent = 0.9 (left) and its power spectrum (right). (**d**) An example of additive signal combining fractional Gaussian noise and periodic signal (left) and its power spectrum (right).

In the brain, tissue oxygenation is determined by the balance between the oxygen supplied by the blood and the oxygen consumed by mitochondria. Both of these processes could contribute to fluctuations in oxygenation. In the brain, increases in neural activity are usually accompanied by vasodilation and increased blood flow/volume that leads to increases in oxygenation [22]. The resulting change in oxygenation will involve an interplay of factors, with the increase in blood flow usually, but not always, driving an oxygen increase [20]. The linkage of oxygenation to neural activity is widely used to infer neural activity non-invasively using BOLD fMRI [23], however, there are many examples of neural and vascular signals departing from this relationship [24–29]. Converging evidence from a large body of studies in both rodents and primates have shown that power in the gamma band (nominally 40-100 Hz) of the local field potential (LFP) is most closely related to the vasodilation that leads to increased blood volume and flow [30–36]. Spiking activity has similar correlations to blood volume as gamma-band LFP power [30, 37, 38], while the correlations for other bands of the LFP are much lower [30, 31,34].

There have been speculations that the ultra-slow (< 1 Hz) electrical signals are the neural correlate of brain hemodynamics [39–41], but frequencies below 1 Hz in the LFPs are of a non-neuronal origin [36]. Because the electrical potential of the blood is negative relative to that of the cerebral spinal fluid [42, 43], changes in the blood volume in the brain will generate ultra-slow potentials. The dilation of arterioles (occurs over seconds) and veins (occurs over tens of seconds) in awake animals’ brain [44–46] will generate changes (< 1Hz) in the LFP [47–50]. The non-neuronal origin of <1 Hz electrical signals have been shown with manipulations that dilate or constrict blood vessels independent of changes in neural activity, such as CO_2_ inhalation [48–51], head-tilt and Valsalva maneuver [47]. Additionally, most amplifiers have circuitry setup to reject these very low frequencies [52], so unless the recording setup is specifically designed to measure at DC frequencies, signals < 1Hz are not of a physiological origin.

Collectively, the neural correlate of vasodilation is increases in gamma-band power in the LFP. Though there are many studies investigating the relationship between neural activity and vasodilation, there is a paucity of studies looking simultaneously measuring neural activity and oxygen changes simultaneously [53–56], with only a handful looking in awake animals [8, 20, 57]. Whether 1/f-like dynamics in brain oxygenation are driven by neural activity bears on the interpretation of hemodynamic imaging. Several fMRI studies have suggested that 1/f-like dynamics exist in human BOLD signals [58–61], and the 1/f-like fluctuations in brain hemodynamics have been interpreted as being driven by 1/f-like fluctuations in neural activity [62, 63]. However, recordings of the LFP in both humans [64] and non-human primates [65] do not seem to show 1/f dynamics in band-limited power.

As 1/f-like oxygen fluctuations are found in other tissue [1, 2], their origin may not be neural, and could come from vascular process. Blood flow and arterial diameter show fluctuations in a similar frequency range as oxygen fluctuations [3]. Additionally, as oxygen is carried by red blood cells (RBCs), fluctuations in the flux of RBCs can drive erythrocyte-associated transients (EATs) in oxygen in the tissue [66–82], and fluctuations in flux of these changes in local oxygenation in the cortex [83–86]. Stalls, brief stoppages in blood flow through capillaries happen sporadically and continuously in the cortex due to transient blockage of blood flow by leukocytes [87–92], which are known to greatly increase vascular resistance [93]. These blockages likely drive changes in tissue oxygenation [94] and increased frequency of these stalls has been linked to neurodegenerative disorders [87, 88, 94].

To understand the relationship between neural activity and 1/f-like oxygen tension oscillations in the brain, we used oxygen polarography to directly measure brain tissue oxygenation in different cortical regions and layers in awake mice. We find that in un-anesthetized, head-fixed mice: (1) cortical oxygenation showed 1/f-like power spectra that are similar across cortical regions and layers; (2) the band-limited power of LFP activity did not show 1/f-like power spectra; (3) there was significant coherence and correlation between neural activity and tissue oxygenation, but both were small; (4) silencing neural activity did not stop 1/f-like fluctuations in brain oxygenation; (5) simulations of erythrocyte flow taking into account the statistics of erythrocyte spacing show that the irregular nature of erythrocyte spacing can generate 1/f-like dynamics in tissue oxygenation. Our results suggest that the driver of 1/f-like oxygenation fluctuations is non-neuronal in origin, and could be due to fluctuations in red blood cell flux.

## Results

We measured tissue oxygenation signals and neural activity from the somatosensory and frontal cortices of awake behaving mice head-fixed on a spherical treadmill [20, 24, 30, 95]. We recorded laminar neural activity with linear multisite probes in 6 mice, laminar oxygenation using polarographic electrodes in 37 mice, and simultaneous neural activity, respiration and oxygen measurements in 9 mice. Additional 5 mice were used to measure RBCs spacing in capillaries using two-photon laser scanning microscopy (2PLSM). We reported results for rest period which only includes data from periods of time when the animal was not locomoting, or for all data, which includes periods of locomotion and rest. We did this because un-anaesthetized mice engage in spontaneous movement frequently, and these spontaneous movements are large drivers of neural activity and hemodynamic signals [30, 36, 96–98]. Specifically, cutaneous sensation during locomotion drives large increases in neural activity in the FL/HL region [20, 24, 99, 100]. The increase in neural activity drives localized increases in blood flow, which is not due to systemic factors [20, 101]. Neural and oxygen measurements were made in the frontal cortex (FC) and in the forelimb/hindlimb region of the somatosensory cortex (FL/HL, identified by cytochrome oxidase staining [102]). All power spectra and frequency domain analyses were done using multi-taper techniques [103], which minimize spectral leakage, using the Chronux toolbox (http://chronux.org/). Portions of this dataset have been published previously [20].

### Brain oxygenation shows 1/f-like power spectrum with a band-limited component

We first asked if tissue oxygen concentrations (PtO_2_) in the cortex of awake mice showed 1/f-like power spectra, as has been observed in the cortex of non-human primates [8]. We collected tissue oxygenation in multiple sites in 37 awake behaving mice (**Figure 2**a). The average duration of recording at each site was 37.0 ± 11.6 minutes. Examination of the resting PtO_2_ trace reveals that oxygen levels show slow fluctuations on the time scale of seconds or longer (**Figure 2**b). The power spectra of tissue oxygen signals, when plotted on a log-log axes, was linear, with a band limited component (in the range of 0.1-0.3 Hz) (**Figure 2**c), as seen in non-human primates [8]. The band-limited oscillations cover the frequency band in which spontaneous arterial oscillations are seen *in vivo* when neural activity is blocked [30] and *ex vivo* in cannulated arteries [104–107]. As a control for any non-physiological sources of this signal [108], we measured PtO_2_ in a dead mouse. The power spectrum of PtO_2_ in the dead mouse was essentially flat (**Figure 2**c), characteristic of white noise (**Figure 1**b), and was several orders of magnitude smaller at all frequencies, ruling out a non-physiological origin of these fluctuations.

**Figure 2.**
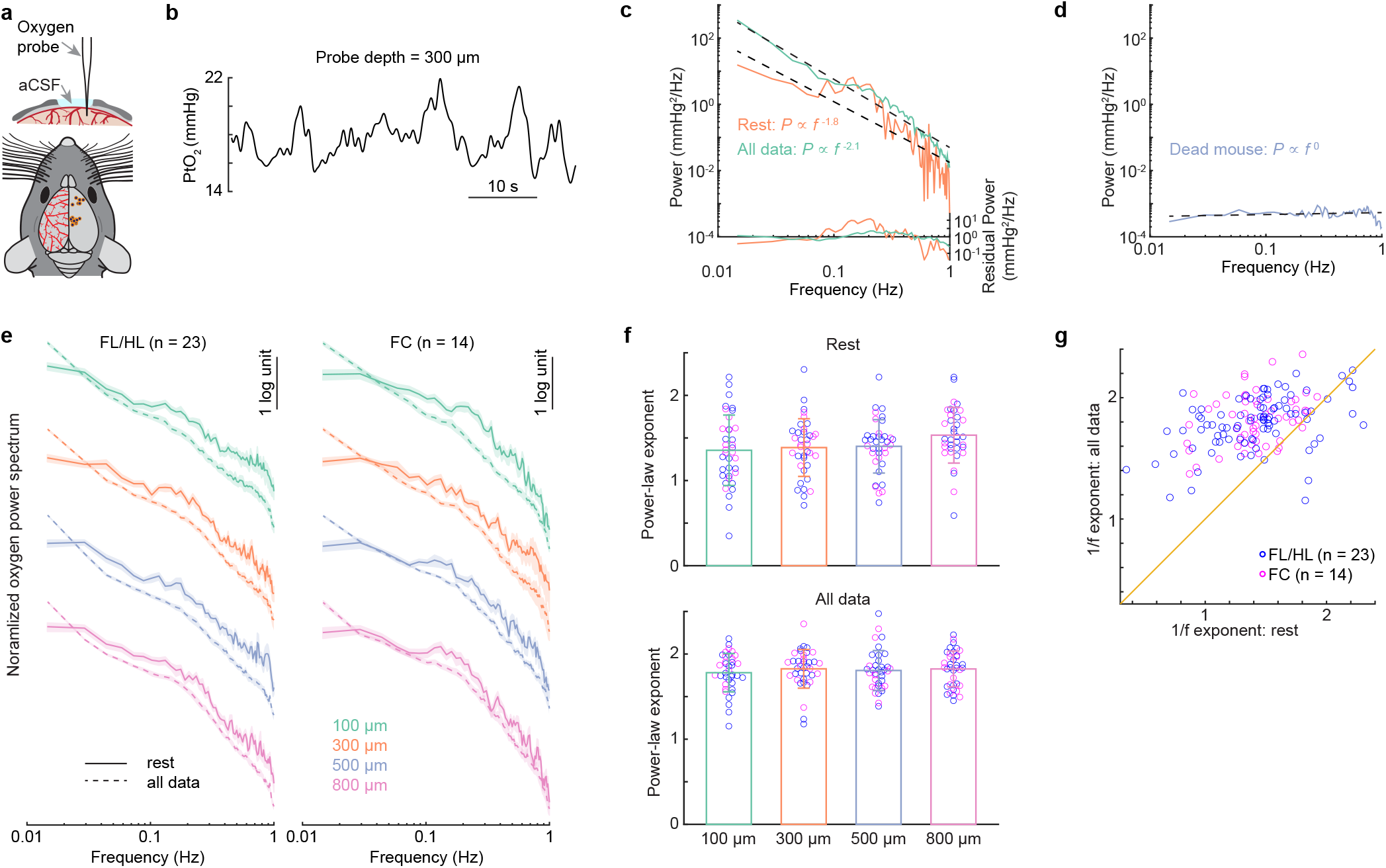
1/f-like dynamics of tissue oxygenation. (**a**) Experimental setup (top) and a summary of measurement sites (bottom). (**b**) Example trace showing the spontaneous fluctuations of tissue oxygenation (PtO_2_) 300 μm below the pia. (**c**) Power spectrum of the PtO_2_ (solid line) as well as its power-law fit (dashed line) using data from resting period (orange) and periods including both rest and locomotion (green). The residual power (i.e., the difference between the power spectrum of the observed oxygen signal and the power-law fit) is shown in the bottom. (**d**) The power spectrum of PtO_2_ (purple) and its linear regression fit (black) from a dead mouse. (**e**) Group average of power spectrum during periods of rest (solid line) and periods including both rest and locomotion (dashed line) across different cortical depths in both FL/HL (n = 23 mice) and FC (n = 14 mice). (**f**) Group average of power-law exponent across different cortical layers during periods of rest (top) and periods including both rest and locomotion (bottom). Blue circles denote the measurements in FL/HL while the magenta circles denote the measurements in FC. (**g**) Scatter plot showing the increase of power-law exponent during periods including locomotion. The orange line indicates the unity line. Data are shown as mean ± SEM in (**e**) and (**f**).

We then qualified the nature of the power spectrum of oxygen fluctuations by fitting it with a power law distribution in the 0.01-1 Hz frequency range. To estimate the power-law exponent, we fitted the oxygen power spectrum using a linear regression (see Methods), to allow comparisons to previous studies [8], though there are caveats to this approach [109]. Averaged across all animals (n = 37), the power law exponent during rest was 1.42 ± 0.21, comparable to what have been observed in un-anaesthetized non-human primates (1.74) using oxygen-sensitive microelectrodes [8], but somewhat larger than those observed in human BOLD studies [58]. We then asked if the exponent of the fit to the power spectrum differed across cortical layers, since there are laminar differences in vascular, mitochondrial, and cellular density [110–112], which could affect the oxygen dynamics [85, 86]. No significant differences were observed in different cortical depths at rest (1.36 ± 0.41 at 100 μm, 1.39 ± 0.34 at 300 μm, 1.40 ± 0.31 at 500 μm, 1.53 ± 0.33 at 800 μm, n = 37 mice, **Figure 2**f, Kruskal-Wallis test, *χ*^2^(3,146) = 5.79, p = 0.1223). We next asked if the 1/f-like dynamics of tissue oxygenation differed between FC and FL/HL, as different power-law exponent has been observed in different brain networks [58]. We did not observe a significant difference between the fitted exponents for FC (n = 14 mice, 1.41 ± 0.13, average across all cortical depths for each animal) and those of the FL/HL (n = 23 mice, 1.42 ± 0.25, average across all cortical depths for each animal, Wilcoxon rank sum test, p = 0.4810). We then asked if the power law fit was affected by behavior, so we fitted the power spectrum of the whole dataset including both rest and locomotion data. Including all the data increased overall power, with most of the power increase occurred at lower frequency (**Supplementary Figure 1**). Including the locomotion periods increased the power law exponent (n = 37 mice, rest: 1.42 ± 0.21, periods including both rest and locomotion: 1.81 ± 0.14, **Figure 2**f, Wilcoxon signed rank test, p <0.0001).

These results show that, just as in primates [8], there are 1/f-like dynamics in the oxygen levels in the cortex of mice. There was also a band-limited component, albeit at a slightly higher frequency than that found in primates [8], close to the vasomotion frequency [30].

### Fluctuations in band-limited power of the LFPs do not show 1/f-like dynamics

To determine whether neural activity exhibited similar dynamics to oxygen signals, we recorded LFPs (1-100 Hz) and multi-unit activity (MUA, 300-3000 Hz) from 16-channel laminar electrodes placed in the FC (n = 4 sites) and FL/HL (n = 6 sites) during wakeful rest and locomotion in a separate group of mice (n = 6 mice, **Figure 3**a). Recording from one site from FC was excluded from this analysis as there was not enough resting data (see Methods). Broadband LFPs showed 1/f-like power spectra above 5 Hz, but not below, and MUA activity have a relatively smaller slope (**Supplementary Figure 2**). The most relevant aspect of the LFP for the oxygen signal is the band-limited power (BLP, **Figure 3**b and c), which is a measure of envelope amplitude changes of LFP oscillations at specific frequency bands. Previous studies have shown that BLP in the gamma-band (40-100 Hz) are best correlated with the time course of vessel dilation and oxygen changes [20, 25, 30–32, 34, 38, 113], we next calculated the power of gamma-band LFP oscillations (see Methods, **Figure 3**d) and estimated the power law fitting exponent. In contrast to the broad-band LFPs, a flat power spectrum was observed in the gamma-band BLP in the frequency range below 1 Hz at rest (−0.15 ± 0.14 at 100 μm, −0.03 ± 0.18 at 300 μm, −0.08 ± 0.16 at 500 μm, 0.11 ± 0.96 at 800 μm, **Figure 3**d left, one-way ANOVA, F(3,35) = 0.42, p = 0.74), which shows a characteristic of white noise (**Figure 1**b). Fitting all the data, including both rest and locomotion periods, significantly increased the power law exponent (rest: −0.04 ± 0.49, run: 0.38 ± 0.55, Wilcoxon rank sum test, p < 0.0001), and showed no laminar difference (0.19 ± 0.37 at 100 μm, 0.31 ± 0.35 at 300 μm, 0.35 ± 0.35 at 500 μm, 0.69 ± 0.89 at 800 μm, Kruskal-Wallis test, *χ^2^* (3, 35) = 2.44, p = 0.4856, **Figure 3**d right). The power spectra of the sub-alpha (1-8 Hz) BLP and beta (10-30 Hz) BLP revealed similar results, characterizing white noise (**Supplementary Figure 3**). This shows that the power spectra of BLP fluctuations of LFPs in mice do not have 1/f-like dynamics, consistent with recordings from non-human primates and humans [64, 65, 114].

**Figure 3.**
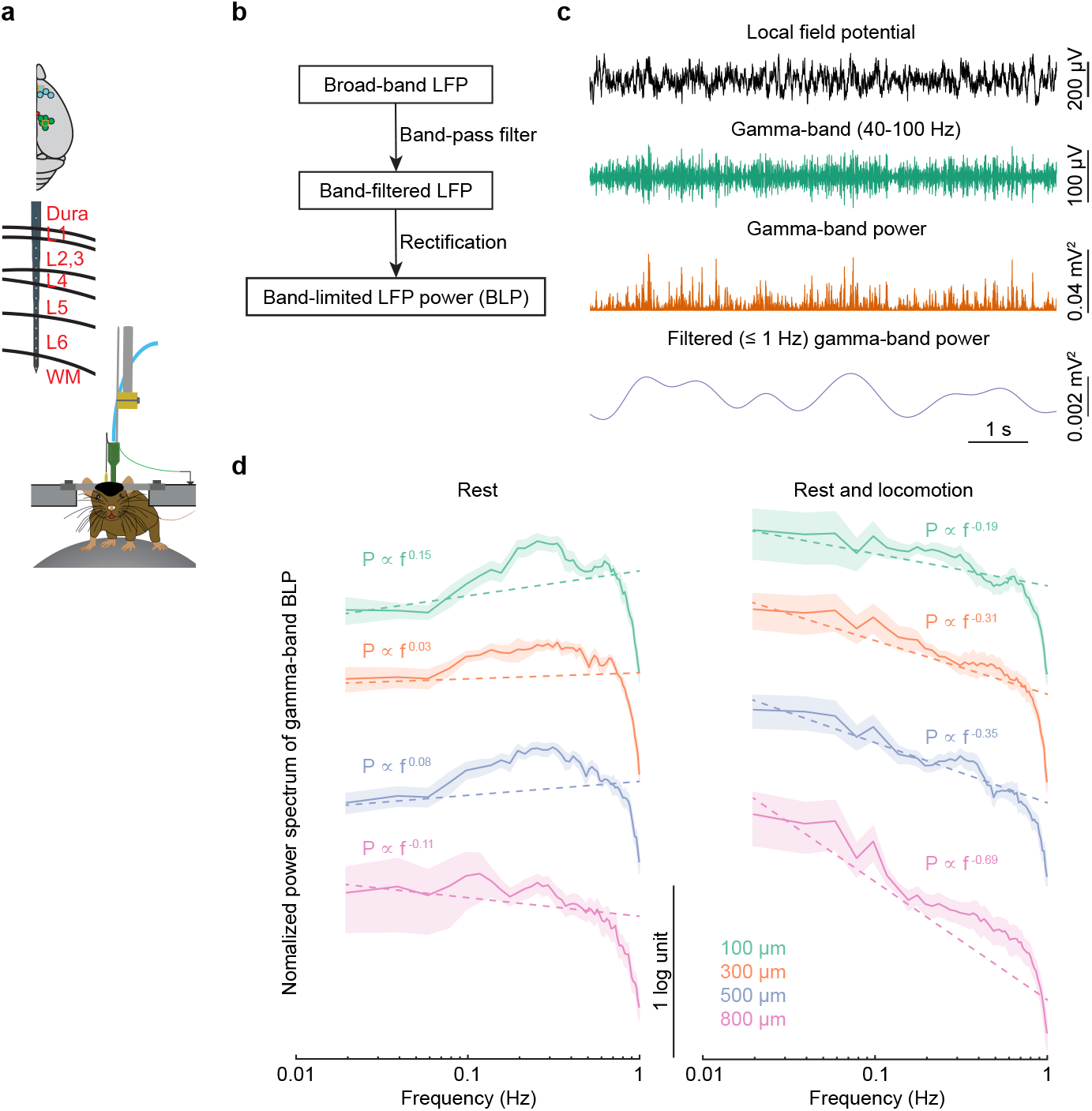
1/f-like dynamics of BLP signals for gamma-band. (**a**) Experimental setup. Top, schematic showing all laminar electrophysiology measurement sites in FC (n = 3 mice) and FL/HL (n = 6 mice). Bottom, schematic showing the layout of the electrodes and measurement depth. (**b**) Schematic outline the general method creating the BLP signals. Raw LFP data was first band-pass filtered and then rectified. The resulting signal was then low pass filtered and resampled. This procedure was applied to each LFP signal for three different frequency ranges. (**c**) Example tracing showing the application of this scheme to one LFP signals, for the gamma-band frequency range. (**d**) Power spectrum of Gamma-band LFP power across cortical depth during rest (left) and periods including rest and locomotion (right). The dashed lines denote the linear regression fit. Data are shown as mean ± SEM.

### Weak correlations between tissue oxygenation dynamics and electrophysiology

The large mismatch between the power-law fit exponents of the power spectrums for band-limited power of LFPs and oxygen fluctuations suggest that their relationship is weak. We then asked how correlated/coherent our oxygen signals were with simultaneously recorded neural activity at rest. To answer this question, we simultaneously measured tissue oxygenation, respiration and LFP activity in 9 animals (see Methods). To differentiate the frequency dependency of the correlation, we calculated the magnitude-squared coherence between oxygen and band-limited power of LFP, as well as the coherence between oxygen and respiratory rate. The magnitude-squared coherence at a given frequency is equivalent to the R^2^ between the two signals bandpass filtered at the frequency [36]. A weak but statistically significant level of coherence between BLP of LFP and oxygen was observed between 0.01-1 Hz, with larger coherence at the lower frequencies (**Figure 4**a-c). However, the magnitude of this squared coherence (which will report the fraction of variance in the signal at that frequency) was low, less than 0.2, implying >80% of the observed variance was not (linearly) predicted by neural activity. This low value should be viewed in light of previous work comparing the correlations of the BOLD signal with simultaneously measured electrophysiological signals in awake primates. These studies found correlation coefficients (R) between gamma-band LFP power and the hemodynamic signals were in the range of 0.3-0.4 [34, 113, 115]. The amount of variance explained (R^2^) by neural activity can be obtained by squaring the correlation coefficient, giving a value in the range of 10-20%, meaning that 80-90% of the observed BOLD signal is uncorrelated with neural activity, similar to our results.

**Figure 4.**
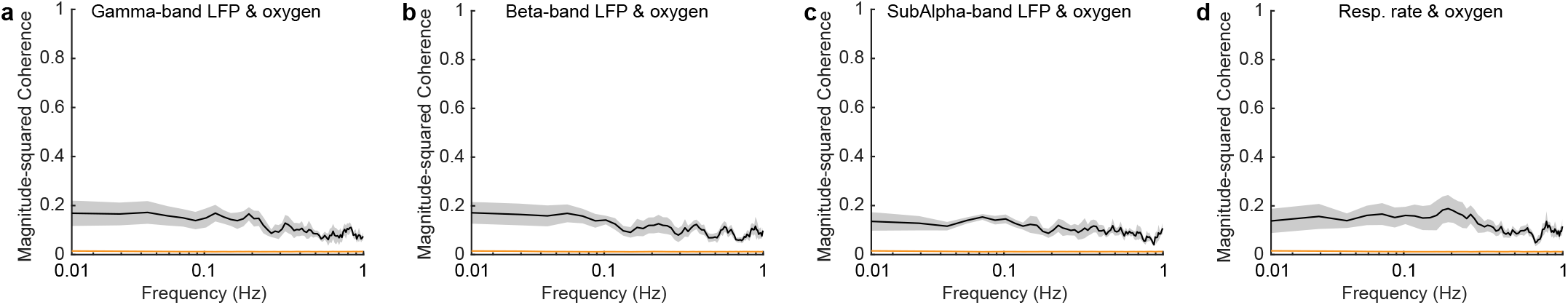
Weak coherence between BLP and tissue oxygenation. Magnitude-squared coherence between gamma-band LFP power (**a**), beta-band LFP power (**b**), sub-alpha band LFP power (**c**), respiratory rate (**d**) and tissue oxygenation during rest. The orange line denotes the 95% confidence interval of the coherence. Data are shown as mean ± SEM.

As another test of how well neural activity can predict changes in oxygenation, we calculated the oxygen hemodynamic response function (HRF, **Figure 5**b) by de-convolving oxygen signals from gamma-band power fluctuations of LFP [20, 30], using the first half of the data from each site. We fit the deconvolved HRF with the sum of two gamma distribution functions (see methods), which is standard in the field [25, 37, 116–119] to create a smoothed HRF. There was a good agreement between the deconvolved and smoothed HRFs (goodness of fit: 0.73 ± 0.39, median ± interquartile range). As increases in gamma-band power will lead to vasodilation and increases in oxygenation, we quantified the positive peak of the gamma distribution fitting. The positive peak of the smoothed HRF (time to peak = 2.99 ± 0.31 s, mean ± SEM, n = 9 mice), and the full-width at half maximum of the HRFs (3.90 ± 3.35 s, median ± interquartile range, n = 9 mice) were comparable to the dynamics of those seen in previous measurements of cerebral oxygen dynamics [120, 121] and BOLD fMRI [122, 123].

**Figure 5.**
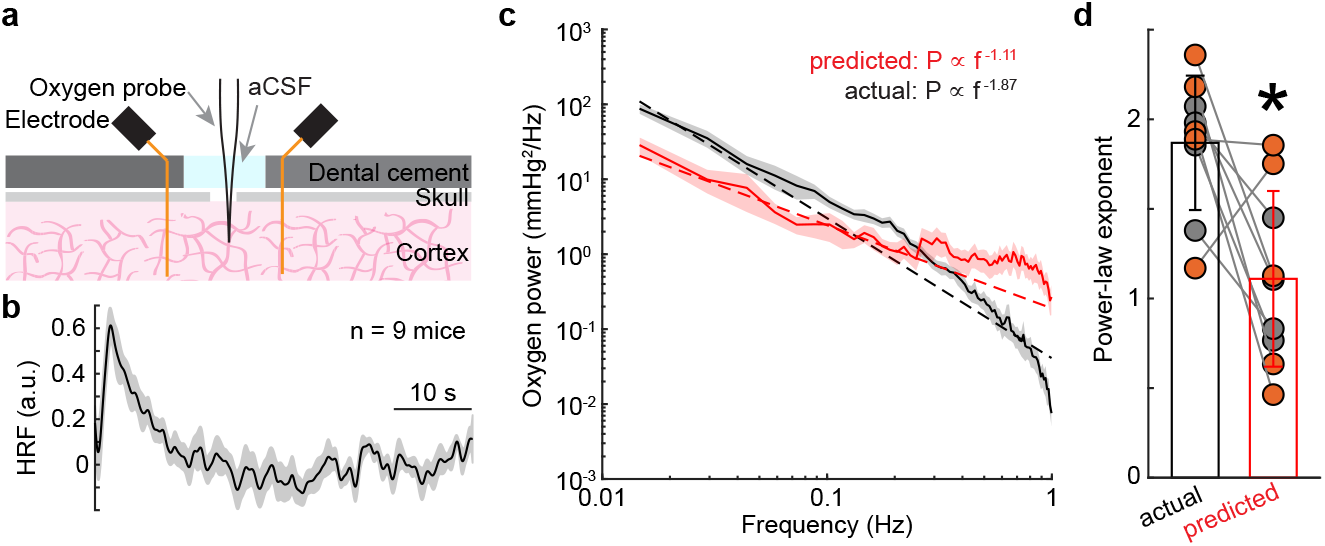
Non-neuronal factors contribute to the 1/f-like dynamics of tissue oxygenation. (**a**) Experimental setup. (**b**) Group average of the hemodynamic response function (HRF) obtained by deconvolving brain tissue oxygenation by gamma-band power of the LFP. Data are shown as mean ± SEM. (**c**) Power spectrum of actual (black) and predicted (red) oxygenation. Data are shown as mean ± SEM. (**d**) Power-law exponent of actual (black) and predicted (red) oxygenation in FL/HL (4 mice, black circle) and FC (5 mice, orange circle). Data are shown as mean ± SD. * paired t-test, t(8) = 3.2520, p = 0.0117.

We then tested how well the HRF to predict tissue oxygenation from neural data. We convolved the HRF with gamma-band power fluctuations using the second half of the data, to get a simulated oxygen signal, which reflects the oxygen component predicted by neural activity (**Supplementary Figure 4**). This model uses the same assumptions built into the analysis of BOLD fMRI data, that the observed signal (oxygen concentration or BOLD) is a linear convolution of the neural activity with an HRF [20, 25, 30, 37, 124]. We then compared the power spectrum between the observed versus the predicted oxygenation. We used data during periods including both rest and locomotion. We found that the oxygen concentration predicted from the neural activity only predicted a small amount of the variance (R^2^) of the signal (R^2^ = 0.04 ± 0.06, n = 9 mice). Furthermore, the power spectrum of the oxygen fluctuations predicted from the neural activity did not show the same frequency dependence as the actual oxygen fluctuations (**Figure 5**c and d, observed: 1.87 ± 0.35, predicted: 1.11 ± 0.46, paired t-test, t(8) = 3.2520, p = 0.0117). Note that the predicted power spectrum was ~63% smaller than the actual power spectrum at frequencies below 0.1 Hz (**Figure 5**c, paired t-test, t(8) = 3.8348, p = 0.0050), indicating that putative non-neuronal components contribute more to the frequencies below 0.1 Hz. These results are consistent with the hypothesis that neural activity does not drive 1/f-like dynamics in tissue oxygenation.

### Impact of suppressing neural activity on tissue oxygenation 1/f-like dynamics

While we found that the majority of the observed oxygen fluctuations could not be explained by neural activity (**Figure 4** and **Figure 5**), if the relationship between neural activity and oxygenation is not captured by the HRF, such as the nonlinearity of brain hemodynamics [125], then some aspect of neural activity might still explain the oxygen fluctuations. To test this possibility mechanistically, we pharmacologically silenced neural activity in the cortex [20, 30], which will also block increases in blood flow mediated by these increases in neural activity [20, 30]. If the 1/f-like power spectrum in oxygen concentration goes away when neural activity is silenced, this would suggest that the fluctuations are due to neural activity and the subsequent vasodilation. If, however, the 1/f-like power spectrum is still present, this would suggest that these oscillations have a non-neuronal origin. To mechanistically understand whether the observed 1/f-like properties of oxygen signals is due to coherent neural activity fluctuations, we superfused the craniotomy with a cocktail of CNQX/AP5/muscimol to suppress neural activity, and recorded tissue oxygenation and LFP simultaneously (**Figure 6**a, **Supplementary Figure 5**). Application of CNQX/AP5/muscimol significantly and substantially suppressed the gamma-band LFP power by 89±8% (Wilcoxon signed-rank test, p = 0.0039) and variance by 77±21% (paired t-test, t(8) = 5.0246, p = 0.0010), but did not change the variance of the tissue oxygenation signal (**Figure 6**b and c, paired t-test, t(8)= 0.7542, p = 0.4723), which suggests that the magnitude of the fluctuations were not reduced by silencing neural activity.

**Figure 6.**
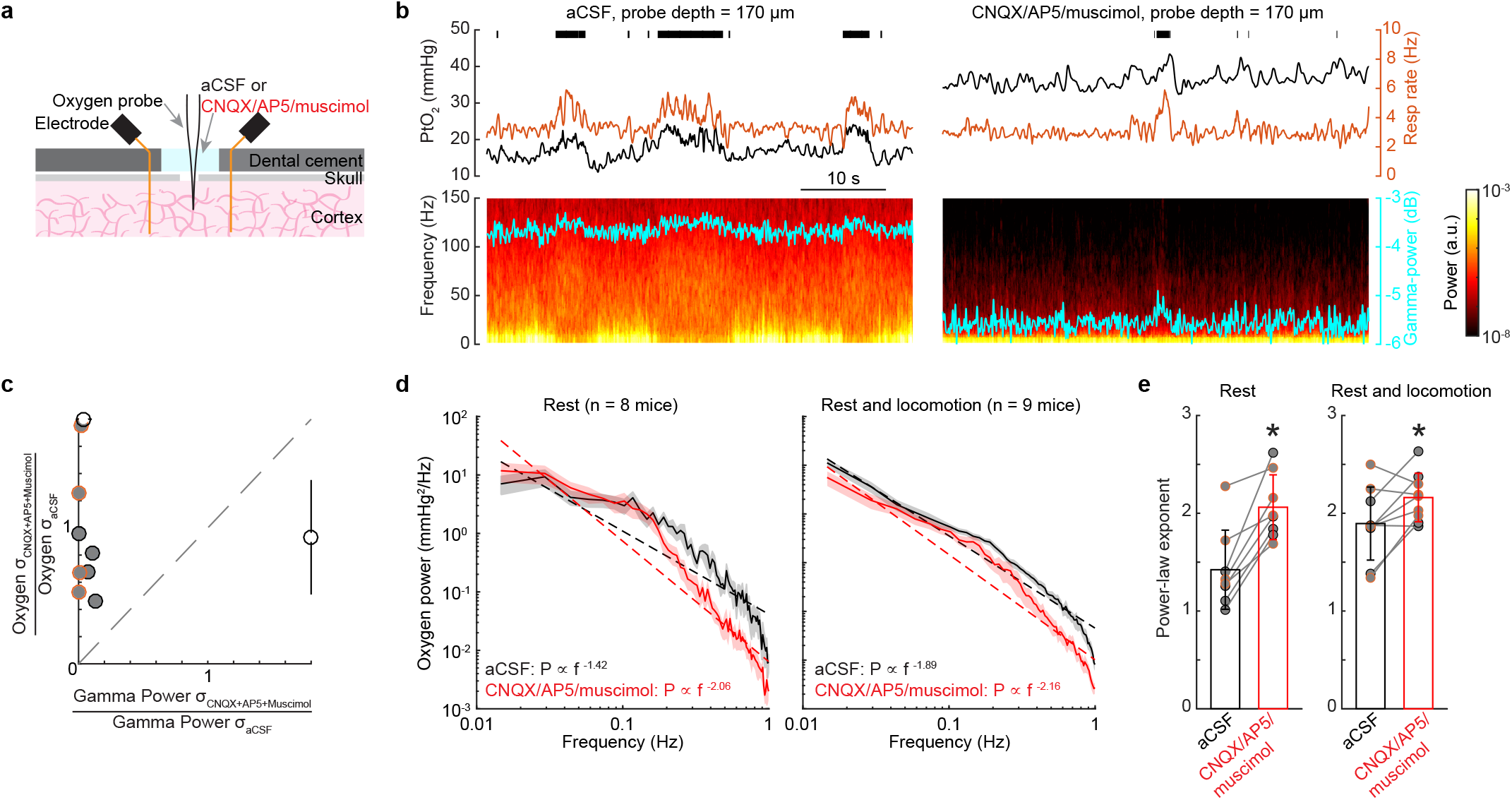
1/f-like fluctuations persist when neural activity is suppressed. (**a**) Experimental setup. (**b**) Example traces showing PtO_2_ responses to locomotion at sites 170 μm below brain surface before (left) and after (right) CNQX/AP5/muscimol. Top, black ticks denote locomotion events; middle, PtO_2_ (black) and respiratory rate (orange) responses to locomotion; bottom, example of data showing spectrogram of LFP (cyan trace showing the gamma-band power). (**c**) Comparison of resting gamma-band power of LFP and PtO_2_ fluctuations quantified with the standard deviation of the signal. Black circles and bars outside the axes show the population mean and SD. The dashed gray line is the unity line. Clustering of the points in the upper left corner shows a pronounced decrease in the neural activity was not accompanied by a decrease in the amplitude of the oxygen fluctuations. (**d**) Power spectrum of tissue oxygen signal before (black) and after (red) application of CNQX/AP5/muscimol using resting data (left) and data including both rest and locomotion (right). (**e**) Power-law exponent of tissue oxygen signal before (black) and after (red) application of CNQX/AP5/muscimol using resting data (left) and data including both rest and locomotion (right).

We then asked if the suppression of neural activity alters the 1/f-like characteristics of the PtO_2_ power spectrum. If the oxygenation fluctuations are driven by neural activity, decreasing neural activity should reduce the amplitude of the oxygen fluctuations. The power spectrum of spontaneous oxygen fluctuations under aCSF had a power law exponent of 1.42 ± 0.40 (n = 8 mice) during rest. Application of CNQX/AP5/muscimol significantly increased the power-law exponent to 2.06 ± 0.33 (paired t-test, t(7) = 5.3358, p = 0.0011, **Figure 6**d and e) during rest. A significant increase of power law exponent was also observed when using the entire dataset (aCSF: 1.89 ± 0.37; CNQX/AP5/muscimol: 2.16 ± 0.25; n = 9 mice, paired t-test, t(8) = 2.6261, p = 0.0304, **Figure 6**d and e). Silencing neural activity did not affect the amplitude of oxygen fluctuations below 0.1 Hz (aCSF: 5.24 ± 4.18 mmHg^2^/Hz; CNQX/AP5/muscimol: 6.74 ± 5.76 mmHg^2^/Hz; Wilcoxon signed-rank test, p = 0.5469), though there was a decrease in the amplitude of oxygen fluctuations above 0.1 Hz during rest (aCSF: 0.49 ± 0.26 mmHg^2^/Hz; CNQX/AP5/muscimol: 0.29 ± 0.27 mmHg^2^/Hz; Wilcoxon signed-rank test, p = 0.0078). These results reflect that the infra-slow (< 0.1 Hz) oscillations in brain oxygenation are not predicted by neural activity, which is consistent with the observation that these oscillations are primarily not driven by neural activity (**Figure 5**c). Taken together, these results show that suppressing neural activity did not abolish 1/f-like oscillations in tissue oxygenation or decrease the amplitude of the oxygen fluctuations, suggesting that non-neuronal contributions are a major driver of these dynamics. Notably, suppressing neural activity does not change the 1/f-like dynamics in both broadband LFP and BLP fluctuations (**Supplementary Figure 5**).

Respiration is a major factor affects brain oxygenation [20], and fluctuations in respiration rate are known to drive substantial changes in BOLD fMRI signals [13, 15, 19, 126]. If the respiration rate shows 1/f-like dynamics [127], this could account for the fluctuations in oxygenation that we see in the tissue when neural activity was suppressed. We found no evidence for 1/f-like dynamics in the respiration rate during rest (**Supplementary Figure 6**). Suppressing neural activity significantly affected the mouse behavior (i.e., reduced the time mice spend locomoting), but did not affect the respiration dynamics during rest (n = 8 mice; fitted exponent for aCSF: 0.27 ± 0.13; CNQX/AP5/muscimol: 0.27 ± 0.04; Wilcoxon signed-rank test, p = 0.8785). Including data during locomotion did not yield an exponent above 1 (n = 9 mice; aCSF: 0.60 ± 0.19; CNQX/AP5/muscimol: 0.46 ± 0.21; Wilcoxon signed-rank test, p = 0.1359). The lack of 1/f-like dynamics in respiration indicate that fluctuations in blood oxygenation due to the fluctuations in respiration are not the origin of the observed 1/f-like oxygen dynamics in the cortex.

### Role of RBCs spacing variations in generating 1/f-like tissue oxygenation fluctuations

The vast majority of oxygen in the blood is carried by RBCs, with the plasma carrying a small fraction of the total oxygen [78, 83, 85, 86], which means heterogeneities in RBCs densities will cause changes in local oxygen supply [67, 70, 72, 78, 83–86]. It has long been appreciated from theoretical models that the tissue oxygenation can vary with the passage of a single RBC, creating an erythrocyte-associated transient (EAT) in tissue oxygenation [66–82]. Recent high-resolution measurements of oxygenation with phosphorescent dyes have confirmed the existence of these transients [67, 72, 83-86], but these measurements require aligning the signals to the passage of the RBCs and would not be able to assay any slow oxygenation change that drive 1/f-like dynamics. Notably, oxygen sensitive electrodes used in the present study lack the temporal resolution to detect the EATs. As RBCs transit through the capillaries in single file flow, the tissue oxygenation outside the capillary fluctuates with their density (**Figure 7**e). Interestingly, there are infrequent “stalls” in RBCs flow in capillaries, caused by leukocytes transiently blocking flow [87, 88, 92, 128, 129]. During a stall, a large RBC spacing will result in a sudden drop of tissue oxygenation within ~15 μm [110] of the capillary. As theoretical work has shown that pulsatile time series generate 1/f-like spectra [130], we asked if the variations in RBCs density through single capillaries had 1/f-like dynamics.

**Figure 7.**
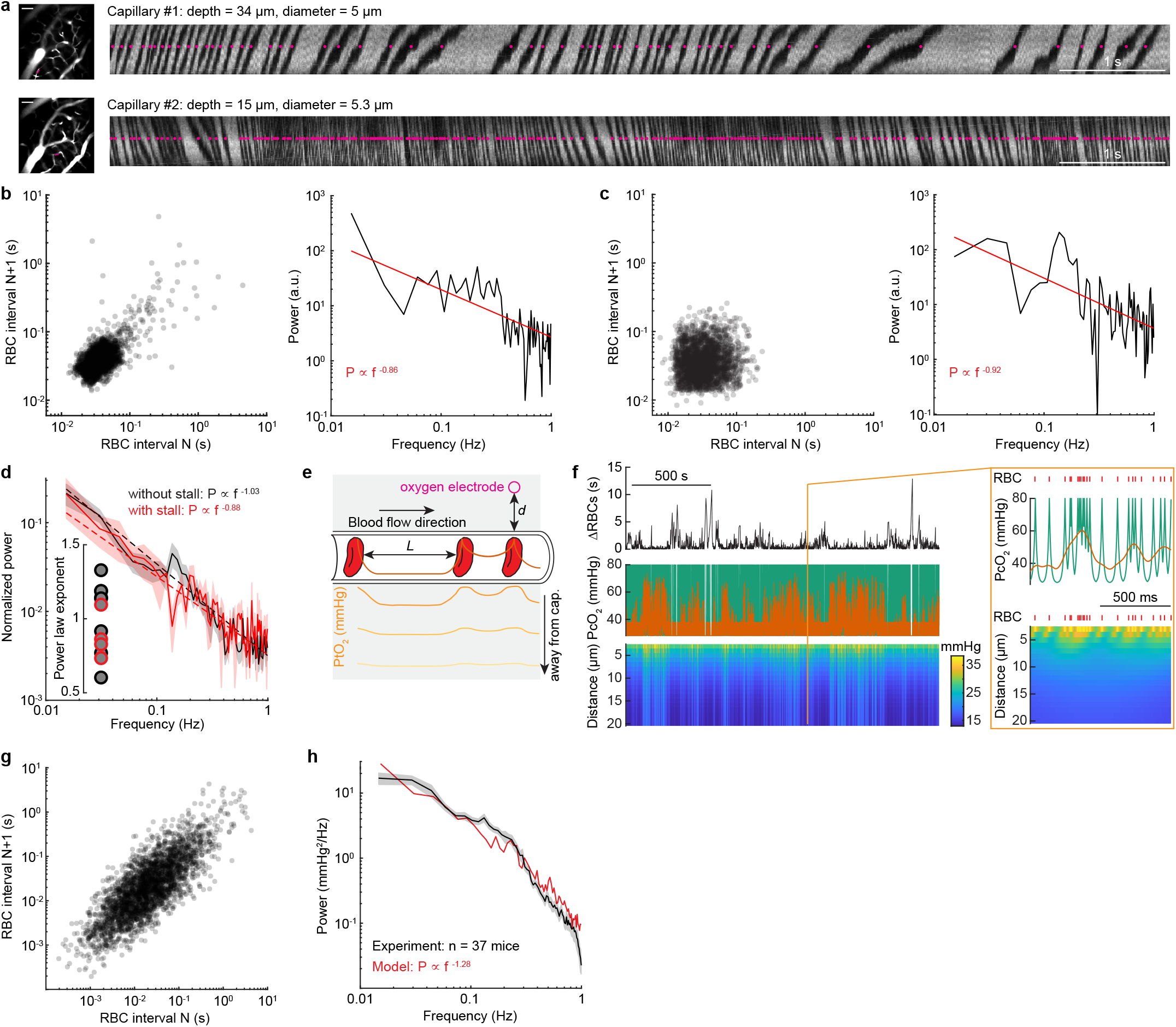
RBCs spacing heterogeneity contributes to 1/f-like oxygen fluctuations. (**a**) Example line scan images showing the RBCs spacing in two different capillaries with (top) and without (bottom) stall events in an example mouse during rest. The images on the left showing the vasculature around the measurement sites, as indicated by the magenta line. Scale bar = 50 μm. The magenta dots indicate the detected RBCs locations. (**b**) Left, lagged scatter plot showing the relationship between consecutive RBC spacing intervals (i.e., autocorrelation trend) of capillary #1. The dark area indicates the density of the RBC spacing. Right, power spectrum for the RBC spacing of capillary #1. The scatter plot showing 3341 RBCs in a 142.8 second resting period. (**c**) As (**b**) but for capillary #2. The scatter plot showing 2603 RBCs in a 100.4 second resting period. (**d**) Power spectrum of inter-RBC transit time for data without stall events (black) and with stall events (red). Inset, fitted power-law exponent for each of the capillaries with (red circle) and without (black circle) stall events. (**e**) Schematic showing that PO_2_ measured at the RBC border decreases with distance and reaches its lowest value between two RBCs. Orange line inside the capillary denotes the oxygenation carried by the RBC and the diffusion to plasma due to intravascular resistance. The gray shaded area denotes brain tissue. Solid traces inside the gray shaded area denotes PtO_2_ at different distance away from the capillary wall (as indicated by the arrow). (**f**) Simulated data showing the fluctuations of RBC spacing (top), oxygenation in the capillary (middle) and oxygen in the tissue (bottom) generated by simulating a Krogh cylinder of 20 μm radius supplied by a capillary of 3 μm radius. The orange box denotes a 1 second segment of the dataset. The red tick marks denote the passage of a single RBC. (**g**) Lagged scatter plot showing the relationship between consecutive RBC spacing intervals for a segment (~3000 RBCs interval) of the simulated data shown in (**f**). (**h**) Comparison of power spectrum of tissue oxygenation measured using polarographic electrodes (black) and generated using the simple model (red) as shown in (**g**). Shaded area denotes 1 SEM.

To answer this question, we measured inter-RBCs spacing in capillaries using 2PLSM to perform line scans along individual capillaries [131]. The plasma is labeled with a fluorescent dye, and RBCs appear as dark streaks (**Figure 7**a), and the pattern of RBCs and plasma was thresholded and binarized to generate a train of point processes. A “stall” event was defined as an inter-RBC spacing of > 1 s. We first quantified the nature of the power spectrum of RBCs arrival fluctuations during rest by fitting the power spectrum of the binarized data (0: plasma; 1: RBC) with a power law distribution in the 0.01-1 Hz frequency range. Binarization makes sense, as the oxygen levels will be high as an RBC passes by, and low when there is only plasma present. We observed 1/f-like dynamics of RBCs spacing, with the exponent range from 0.6 to 1.3 (0.98 ± 0.23, 12 capillaries in 5 mice, **Figure 7**b-d). Note that the occurrence of RBCs “stall” events did not significantly affect the fitted exponent (**Figure 7**d, two sample t-test, p = 0.2848; 0.88 ± 0.15, 4 capillaries with stall events; 1.03 ± 0.25, 8 capillaries without stall events). The inter-RBC interval correlations show that RBCs tend to cluster together in time (**Figure 7**b). The rare occurrence of “stall” events in our data is consistent with previous work [92, 94, 128].

We then developed a simple computational model to determine how the delivery of oxygen by RBCs is affected by the statistics of RBC passages. Given the volume of tissue sampled by the electrode [132], and the spacing of capillaries [110], the oxygen signal at our electrode will be dominated by the nearest capillary. We generated a time series of RBCs spacing (see Methods) utilizing data from our 2PLSM observations and data from previous studies [92, 128], as well as the previously measured oxygen changes associated with RBC passage [83–86]. We only considered the tissue oxygenation transients caused by the RBCs spacing. The RBCs (with high PO2) and plasma gaps (with relatively low PO2) alternately passed through the capillary, and the EAT maybe visualized by considering a fixed measurement site on the capillary wall (**Figure 7**e). As the tissue response time is much slower compared to the RBC transit time (due to the low-pass filtering nature of the oxygen diffusion dynamics), the oxygen delivery from capillaries decays rapidly with distance at higher temporal frequencies. This means that the tissue oxygenation will be a smoothed version of the EATs (**Figure 7**e). Based on our observation, we created a RBC time series with a power spectrum with a fitted exponent 1. The RBCs were considered as a point process, and the EATs surrounding each RBC were modelled using an exponential decay (**Figure 7**e), based on the data measured using two-photon phosphorescent imaging [83–86]. The tissue response time [73] was also considered when calculating the tissue oxygen at different length from the capillary, as oxygen delivery from capillaries decays rapidly with distance at high frequencies of pulsatile flow in the vessels (**Figure 7**e). We generated simulations of oxygenation of comparable length of our data using polarographic electrodes (~40 minutes), and examined the power spectrum. **Figure 7**f illustrated a representative RBC train we modelled with 5% stall events and a power spectrum with a fitted exponent 1. The generated data has shown that RBC spacing fluctuations have long-range autocorrelations (**Figure 7**g), as seen in the data. We assumed that oxygenation of each RBC (PO_2_ = 80 mmHg) and oxygenation in each plasma gap (PO_2_interRBC = 28 mmHg) were constant, giving a EAT magnitude of 52 mmHg, consistent with measurements in awake animals [85, 86]. Altering the EAT magnitude did not affect the slope (**Supplementary Figure 7**). The power spectrum of tissue oxygenation fluctuations originated from RBCs heterogeneity had a slope 1.28, and the simulations showed very close agreement with the data (**Figure 7**h).

Taken together, our simulations show that variations of RBC spacing alone can generate 1/f-like oxygen fluctuations in the capillaries and surrounding tissue. This simple model points to an intriguing possibility that the discrete nature of RBCs may play a more important role in 1/f-like dynamics of oxygen supply. As the fluctuations of RBCs are largely attributed to non-neuronal mechanisms [129, 133], suggesting a non-neuronal origin of 1/f-like oscillations in tissue oxygenation.

## Discussions

We found that oxygen dynamics in the mouse cortex show large, low frequency oscillations which were absent in the band limited power of the LFP. These fluctuations were present in all cortical layers and multiple regions both when the mouse was at rest and during behavior. These fluctuations were weakly correlated with neural activity and persisted when neural activity was pharmacologically suppressed. Simulations based on physiological measurements showed that the stochastic, correlated fluctuations in the number of red blood cells could account driving these dynamics.

A large vascular contribution to the 1/f-like dynamics in tissue oxygen could potentially explain many disparate observations. It would explain why 1/f-like dynamics are seen in tissue oxygenations throughout the body [1, 5, 8, 9], why we see similar oxygen dynamics across layers and cortical regions (**Figure 2**) even though there are large differences in neural activity and vascular density across regions and layers [110–112]. Fluctuations in oxygenation generated by the stochastic passage of RBCs are ‘noise’ and could explain the low correlations and coherences between oxygen and neural activity observed both in our experiments and in BOLD fMRI measures [34, 113, 115].

Besides the RBC heterogeneity, there may be other contributors to oxygen dynamics. The tissue oxygenation is determined by an interplay of metabolism and supply. Fluctuations in the consumption of oxygen by mitochondria may also contribute to the 1/f-like dynamics both at the single cell level [134–137] and at the system level [138]. Moreover, measurements in awake and sleeping cats have shown that existence of spontaneous oscillating metabolic phenomenon in cortex that is not directly related to neural activity [139]. However, oxygen consumption by neurons in rat hippocampal slices is closely tied to neural activity [140] and oxygen levels in the slices lack the fluctuations seen in perfused tissue. This suggests that non-neural related metabolic activity is a potential contributor to the observed oxygen 1/f-like dynamics.

In addition to stalls in flow caused by vessel occlusion by leukocytes [87, 88, 92, 94, 128, 129], the very nature of the RBC-plasma suspension will drive low-frequency fluctuations in flow and RBC heterogeneity [133, 141]. RBCs have different rheological properties than the plasma [142, 143], making the Newtonian descriptions of fluid flow that works for larger vessels inapplicable to the flow through capillaries. Because of the higher effective viscosity of RBC than the plasma, stochastic fluctuations in the number of RBC in a vessel changes the effective resistance of the vessel, which can lead to low frequency fluctuations in RBC flow [133, 141]. Besides these coagulation factors affecting capillary blood flow, the temporal and spatial capillary flow dynamics are also affected by the contractile dynamics of pericytes [144–146]. Though we modelled the RBC and oxygenation dynamics in one single capillary, the vasculature topology will introduce additional level of complexity, and affects how the RBCs are distributed within the network [133, 141] and subsequently affect oxygenation dynamics.

Collectively, our results suggest that processes affect the location and frequency of stalling events and RBC heterogeneity will greatly impact the levels of oxygen in the tissues. Besides the aforementioned leukocytes caused “stall” events [87, 88, 92, 94, 128, 129], the temporal dynamics of RBCs in the capillary may also reflect general features of the capillary network supply the same brain region [133, 141]. It is also possible that the arterial blood pressure fluctuations [147], heart beat dynamics [148], upstream vessel diameter oscillations [44], rheological properties of flowing blood [142, 143] and anatomical structures of capillary [149], and the dilation of capillaries and activity of pericytes[144, 146, 150, 151] contribute to the stochastic RBC dynamics.

## Acknowledgements

We thank Nanyin Zhang, Yuncong Ma for comments on this manuscript. This work was supported by the National Institutes of Health grant R01NS078168 to P.J.D., and the National Institutes of Health grant R01NS108407 “Understanding cellular architecture of the neurovascular unit and its function in the whole mouse brain” to Dr. Yongsoo Kim (Department of Neural and Behavior Sciences, College of Medicine, The Pennsylvania State University).

## Author contributions

Q.Z. and P.J.D. designed the project. Q.Z. performed experiments using polarographic oxygen electrodes, pharmacological manipulation, and two-photon laser scanning microscopy. K.W.G. performed experiments measuring laminar electrophysiology. Q.Z. performed all the data analysis and implemented the computational modelling. Q.Z. and P.J.D. wrote the manuscript. P.J.D. supervised experiments, data analysis, and preparation of the manuscript.

## Materials and Methods

Portions of the data used in this study have been published previously [20]. This study was performed in strict accordance with the recommendations in the Guide for the Care and Use of Laboratory Animals of the National Institutes of Health. All procedures were performed in accordance with protocols approved by the Institutional Animal Care and Use Committee (IACUC) of the Pennsylvania State University (protocol #201042827).

### Animals

A total of 47 C57BL/6J mice (3-8 months old, 25-35 g, Jackson Laboratory) were used. Recordings of laminar cortical tissue oxygenation were made from 37 mice [23 (13 male and 10 female) in the somatosensory cortex (FL/HL) and 14 (7 male and 7 female) in the frontal cortex (FC)] using Clark-type polarographic microelectrode. Simultaneous measurements of cortical tissue oxygenation using polarographic electrodes, respiration and local field potential were conducted in 9 mice [5 (4 male and 1 female) in FL/HL and 4 (2 male and 2 female) in FC]. Six of these mice were also used for laminar cortical tissue oxygenation measurements. Local field potential and spiking activity of different cortical layers were measured using laminar electrodes in a separate set of 7 male mice (4 in FC and 6 in FL/HL, 3 mice were measured in both FL/HL and FC simultaneously). Two-photon laser scanning microscopy (2PLSM) imaging was conducted in 5 mice (12 capillaries, 3 male and 2 female, in FL/HL). Mice were given food and water *ad libitum* and maintained on 12-hour (7:00–19:00) light/dark cycles. All experiments were conducted during the light period of the cycle.

### Surgery

All surgeries were performed under isoflurane anesthesia (in oxygen, 5% for induction and 1.5-2% for maintenance). A custom-machined titanium head bolt was attached to the skull with cyanoacrylate glue (#32002, Vibra-tite). The head bolt was positioned along the midline and just posterior to the lambda cranial suture. Two self-tapping 3/32” #000 screws (J.I. Morris) were implanted into the skull contralateral to the measurement sites over the frontal lobe and parietal lobe. A stainless-steel wire (#792800, A-M Systems) was wrapped around the screw implanted in the frontal bone for use as an electrical ground for cortical tissue oxygenation and neural recordings. For capillary blood flow velocity measurements using 2PLSM (n = 5 mice), a polished and reinforced thin-skull (PoRTS) window was made covering the right hemisphere as described previously [20, 24, 30, 45, 101, 152]. For simultaneous measurement of tissue oxygenation and neural activity (n = 9 mice), we implanted two electrodes to measure LFP signals differentially. Electrodes were made from Teflon-coated tungsten wire (#795500, A-M Systems) with ~1 mm insulation striped around the tip. The electrodes were inserted into the cortex to a depth of 800 μm at 45° angle along the rostral/caudal axis using a micromanipulator (MP-285, Sutter Instrument) through two small burr holes made in the skull. The two holes for the electrodes were made ~1-1.5 mm apart to allow insertion of the oxygen probe between the two electrodes in following experiments. The holes were then sealed with cyanoacrylate glue. Following the surgery, mice were then returned to their home cage for recovery for at least one week, and then started habituation on experimental apparatus. Habituation sessions were performed 2-4 times per day over the course of one week, with the duration increasing from 5 min to 45 min.

### Physiological measurements

Data from all experiments, except experiments using 2PLSM, were collected using custom software written in LabVIEW (version 2014, National Instruments).

#### Behavioral measurement

The treadmill movements were used to quantify the locomotion events of the mouse. The animal was also monitored using a webcam (Microsoft LifeCam Cinema®) as an additional behavioral measurement.

#### Cerebral tissue oxygenation measurement using polarographic electrode

On the day of measurement, the mouse was anesthetized with isoflurane (5% for induction and 2% for maintenance) for a short surgical procedure (~20 min). A small (~100 x 100 μm) craniotomy was made over the frontal cortex (1.0 to 3.0 mm rostral and 1.0 to 2.5 mm lateral from bregma) or the forelimb/hindlimb representation in the somatosensory cortex (0.5 to 1.0 mm caudal and 1.0 to 2.5 mm lateral from bregma), and dura was carefully removed. The craniotomy was kept moist with warm artificial cerebrospinal fluid (aCSF) and porcine gelatin (Vetspon). The mouse was then moved to and head-fixed upon the spherical treadmill. Oxygen measurements started at least one hour after cessation of anesthesia to minimize the effects of anesthesia [85, 95, 153].

Cerebral tissue oxygenation was recorded with a Clark-type oxygen microelectrode (OX-10, Unisense A/S, Aarhus, Denmark). A total of 9 oxygen electrodes were used in this study, with an average response time of 0.33 ± 0.11 seconds (n = 9 probes). No compensation for the delay was performed. The oxygen electrodes were calibrated in air-saturated 0.9% sodium chloride (at 37°C) and oxygen-free standard solution [0.1M sodium hydroxide (SX0607H-6, Sigma-Aldrich) and 0.1M sodium ascorbate (A7631, Sigma-Aldrich) in 0.9% sodium chloride] before and after each experiment. The linear drift of the oxygen electrode signal (1.86±1.19% per hour) was corrected by linearly interpolating between pre- and post-experiment calibrations. The oxygen electrode was connected to a high-impedance picoammeter (OXYMeter, Unisense A/S, Aarhus, Denmark), whose output signals were digitalized at 1000 Hz (PCI-6259, National Instruments). Current recordings were transformed to millimeters of mercury (mmHg) using the calibrations with air-saturated and oxygen-free solutions.

Oxygen electrodes allow long-duration, quantitative measurements of the average oxygen tension from a small volume (~20 μm radius) of parenchymal tissue. And the stability of the electrode provides long duration measurements which are required to estimating the power at ultra-low frequencies. For oxygen polarography measurements, the oxygen microelectrode was positioned perpendicular to the brain surface and advanced into the cortex with a micromanipulator (MP-285, Sutter Instrument). Measurement site was chosen to avoid large pial vessels. The depth zero was defined as when the tip of the oxygen microelectrode touches the brain surface under visual inspection. The probe was then advanced to depth of 100, 300, 500 and 800 μm below the pia, and 30-40 minutes data were recorded for each depth. After advancing the electrode, we waited at least 5 minute before the start of each recording.

In experiments investigating effects of suppressing neural activity on cortical tissue oxygenation dynamics, a cocktail of ionotropic glutamate receptor antagonists 6-cyano-7-nitroquinoxaline-2,3-dione (CNQX, 0.6 mM), NMDA receptor antagonist (2R)-amino-5-phosphonopentanoic acid (AP5, 2.5 mM) and GABAA receptor agonist muscimol (10 mM) were applied to suppress neural activity. All drugs were applied topically over the craniotomy and were allowed to diffuse into the cortical tissue for at least 90 min before the oxygen measurements. The efficacy of the CNQX/AP5/muscimol cocktail was validated with simultaneously recorded neural activity. Neural data were amplified 1000x and filtered (0.1 – 10k Hz, DAM80, World Precision Instruments) and then sampled at 30k Hz (PCI-6259, National Instruments). The oxygen signal in these experiments was recorded at a depth of ~100-200 μm.

In experiments investigating effects of suppressing neural activity on cortical tissue oxygenation dynamics, respiration was also simultaneously recorded. Measurements of breathing were taken using 40-guage K-type thermocouples (TC-TT-K-40-36, Omega Engineering) placed near the mouse’s nose (~ 1 mm), with care taken to not contact the whiskers. Data were amplified 2000x, filtered below 30 Hz (Model 440, Brownlee Precision), and sampled at 1000 Hz (PCI-6259, National Instruments). Downward and upward deflections in respiration recordings correspond to inspiratory and expiratory phases of the respiratory cycle, respectively. We identified the time of each expiratory peak in the entire record as the zero-crossing point of the first derivative of the thermocouple signal.

At the end of the experiment, the mouse was deeply anesthetized, and a fiduciary mark was made by advancing an electrode (0.005” stainless steel wire, catalog #794800, A-M systems) into the brain with a micro-manipulator to mark the oxygen measurement site.

#### Laminar electrophysiology

Laminar electrophysiology recordings were performed in a separate set of mice (n = 6). On the day of measurement, the mouse was anesthetized using isoflurane (in oxygen, 5% for induction and 2% for maintenance). Two small (1×1 mm^2^) craniotomies were performed over the frontal cortex (1.0 to 2.5 mm rostral and 1.0 to 2.5 mm lateral from bregma) and FL/HL representation in the somatosensory cortex (0.5 to 1.0 mm caudal and 1.0 to 2.5 mm lateral from bregma) over the contralateral hemisphere, and the dura was carefully removed. The craniotomies were then moistened with warm saline and porcine gelatin (Vetspon). After this short surgical procedure (~20 minutes), the mouse was then transferred to the treadmill where it was head-fixed. Measurements started at least one hour after the cessation of anesthesia [95, 153].

Neural activity signals were recorded using two linear microelectrode arrays (A1×16-3mm-100-703-A16, NeuroNexus Technologies). The electrode array consisted of a single shank with 16 individual electrodes with 100 μm inter-electrode spacing. The signals were digitalized and streamed to SmartBox™ via a SmartLink headstage (NeuroNexus Technologies). The arrays were positioned perpendicular to the cortical surface, one was in the FL/HL and the other one was in the FC on the contralateral side. Recording depth was inferred from manipulator (MP-285, Sutter Instrument) recordings. The neural signals were filtered (0.1-10k Hz bandpass), sampled at 20k Hz using SmartBox 2.0 software (NeuroNexus Technologies).

#### Measuring RBC spacing in capillaries using 2PLSM

Two-photon imaging was performed with a Sutter Moveable Objective Microscope. A MaiTai HP (Spectra-Physics, Santa Clara, CA) laser tuned to 800 μm was used for fluorophore excitation. Before imaging, the mouse was briefly anesthetized with isoflurane (5% in oxygen), retro-orbitally injected with 50 μL of 70 kDa fluorescein-conjugated dextran (Sigma-Aldrich) prepared at a concentration of 5% (weight/volume) in sterile saline to label plasma, and then fixed on a spherical treadmill. Imaging was done with a 20X, 1.0 NA objective (Olympus, XLUMPFLN). Control of 2PLSM and data acquisition was accomplished using MScan software (Sutter Instruments). All imaging with the water-immersion lens was done with room temperature distilled water. Wide-field images were collected to generate vascular maps of the entire window for navigational purposes. High-resolution images of the vasculature were collected using a 500 by 500 μm field for measurement of capillary diameter. Capillary diameter was measured using ImageJ. To measure RBC velocity and RBC spacing, line scan images were collected from individual capillaries. RBCs appeared as tilted dark shadows on a bright background due to the fluorescein-conjugated dextran contained in the blood plasma (**Figure 7**a), and these shadows were counted.

### Data analysis

All data analyses were performed in Matlab (R2015b, MathWorks) using custom code.

#### Locomotion events identification

Locomotion events [20, 24, 30, 154] from the spherical treadmill were identified by first applying a low-pass filter (10 Hz, 5^th^ order Butterworth) to the velocity signal from the optical rotary encoder, and then the absolute value of acceleration (first derivative of the velocity signal) was thresholded at 3 cm/s^2^. Periods of locomotion were categorized based on the binarized detection of the treadmill acceleration:

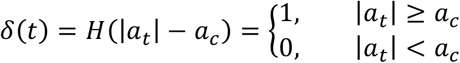

where *a_t_* is the acceleration at time t, and *a_c_* is the treadmill acceleration threshold.

#### Spontaneous activity

To characterize spontaneous (non-locomotion-evoked) activity, we defined “resting” periods as periods at least 4 seconds after the end of previous locomotion event and lasting no less than 60 seconds.

#### Oxygen data preprocessing

Oxygen data from polarographic electrodes were first low-pass filtered (1 Hz, 5^th^ order Butterworth). The oxygen data were then down-sampled to 30 Hz to align with binarized locomotion events.

#### Laminar neural activity

The neural signal was first digital filtered to obtain the local field potential (LFP, 0.1-300 Hz, 5^th^ order Butterworth) and multiunit activity (MUA, 300-3000 Hz, 5^th^ order Butterworth) [20, 24, 30]. Time-frequency analysis of LFP signal was conducted using multi-taper techniques (Chronux toolbox version 2.11, http://chronux.org/) [103]. The power spectrum was estimated with a 1 second window with ~1 Hz bandwidth averaged over nine tapers. MUA signals were low-pass filtered (5 Hz, Bessel filter). The locomotion-evoked LFP power spectrum was converted into relative power spectrum by normalizing to the 3 second resting period prior to the onset of locomotion. Spike rate was obtained by counting the numbers of events that exceed an amplitude threshold (three standard deviations above background) in each 1 millisecond bin.

To examine raw LFP or BLP modulations at different frequency bands, we first used a third-order Butterworth filter to apply zero-phase bandpass filtering to the raw LFP according to the following frequency bands: sub-alpha, 1-8 Hz; beta, 10-30 Hz; and gamma: 40-100 Hz. The resulting BLP signals were squared and full-wave rectified. They were then resampled to 20 Hz after low-pass filtering below 1 Hz. These steps are illustrated in (**Figure 3**b and c).

The spike train were extracted from each channel of the laminar electrode. Firing-rate signals in these data were smoothed with a Gaussian kernel with FWHM of 10 ms to generate a continuous signal.

#### Magnitude-squared coherence

We used coherence analysis [155] to reveal correlated oscillations and deduce functional coupling among different signals. The magnitude squared ordinary coherence between two signals x and y are defined as

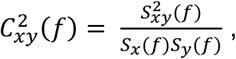

where *S_x_*(*f*) and *S_y_*(*f*) are the auto-spectra of the signals, and *S_xy_*(*f*) is the cross-spectrum.

#### Quantifying the oxygen fluctuations predicted by the neural activity

We considered the neurovascular relationship to be a linear time invariant system [37, 156, 157]. To provide a model-free approach to assess the relationship between tissue oxygenation and neural activity, hemodynamic response function (HRF) was calculated by deconvoluting tissue oxygenation signal to gamma-band LFP power, using the following equation:

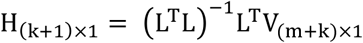

H is the HRF, V is the tissue oxygenation signal. L is a Toeplitz matrix of size (m+k) x (k+1) containing measurements of gamma-band LFP power (n):

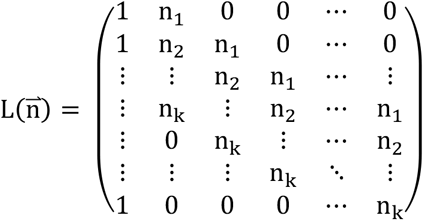

To estimate how much variance of oxygenation the neural activity can predict, we first split the observed data into two segments with equal length. We then calculated the HRF using the first half of the observed data. We smoothed the HRF using a Savitzky-Golay filter (3^rd^ order, 11-point frame length). Next, we convolved the HRF with the gamma-band LFP power from the second half, and estimated the oxygenation predicted by neural activity using the following equation,

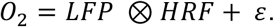

The efficacy of the prediction was quantified by calculating the correlation coefficient (R) between the prediction and actual oxygenation. The process was shown in **Supplementary Figure 4**.

#### Hemodynamic response function kernel fitting

To quantify the temporal features of HRF, the HRF for tissue oxygenation was fitted using two gamma-variate fitting process [25, 37, 116–118] using a gamma-variate function kernel of the following form,

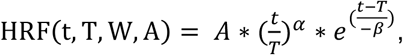

where α = (*T/W*)^2^ * 8.0 * log(2.0), β= *W*^2^/(*T* * 8.0 * log(2.0)). For modeling HRF using a gamma-variate function kernel, we used a downhill simplex algorithm minimizing the sum square difference between measured and predicted hemodynamics. The goodness of fit was quantified as 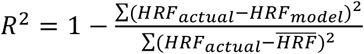, where 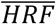 is the mean value of the actual HRF. The amplitude (A), time-to-peak (T) and full-width at half maximum (W) of the kernel was then calculated.

### Modeling RBC spacing effects on tissue oxygenation

We identified the location of each RBC using custom code written in MATLAB from line scan images using 2PLSM (**Figure 7**a). Data was first undergoing visual inspection to determine if the quality was sufficient for reliable RBC detection. To calculate the power spectral density of RBCs train, we estimated using a function specifically for point processes (Chronux toolbox function: mtspectrumsegpb). To estimate the RBC interval distribution, we pooled all observed RBC intervals during rest from different animals (n = 5 mice) together to determine the probability density function (PDF). However, it has been reported that only a small amount of segments experiencing a stall at any given instant in awake mice ([128], 0.5%) and anesthetized mice ([92], 2.1%), which makes the observation of capillaries with a cessation of RBC flow challenging. It is also not practical to measure a large number of capillaries with a sufficiently long duration to characterize the temporal dynamics using 2PLSM, as it requires to focus on a limited number of segments with high magnification and a narrow field of view. To avoid the bias of PDF estimation due to the rare occurrence of “stall” events in our experiments, we also estimated the PDF of “stall” events using data from [128]. Combining these two PDFs, we estimated a new PDF of RBC intervals to generate a synthetic dataset (matlab function: normrnd). As the consecutive RBC intervals are not totally random and has a power law slope close to 1 (**Figure 7**b), we introduced long-range autocorrelation using inverse Fourier transform.

Using the generated RBC train time series, we then simulated the oxygenation change inside the capillary and in the nearby brain tissue (**Figure 7**e and f). As in capillaries, RBCs travel in single file, separated by plasma gaps of variable lengths, therefore, the capillary blood is not a homogenous oxygen source, but has heterogenic dynamics. We therefore assumed that 1) the tissue is primarily oxygenated by the nearest capillary. 2) The space between RBC and capillary wall is minimal, and that the capillary wall does not hinder the transport of oxygen. Therefore, the oxygen concentration profile is continuous between blood and tissue. 3) Oxygen transport within the tissue is assumed to be solely by molecular diffusion and governed by Fick’s second law of diffusion. 4) The rate of consumption of oxygen by the tissue surrounding the capillary is constant. Under these assumptions, over time, the level of oxygen within the tissue rises until the amount of oxygen lost by the passing cells converges to a quasi-steady level. At this quasi-steady state, the oxygen level in the proximity of the capillary fluctuates between a maximum reached just after the passage of a RBC and a minimum midway prior to the arrival of the next RBC (**Figure 7**e and f). The amount of oxygen delivered by a RBC to the tissue slice is the summation of the oxygen mass gained and consumed within the tissue during its residence.

To keep the model tractable, the geometry of the erythrocytes was not considered (for a complete model, see [81]), and the erythrocyte as an oxygen source is kept as a point-like oxygen source [68]. The oxygen tension for each RBC was set to be the same, and the diffusion of oxygen from RBC to plasma was simulated with an exponential decay kernel measured in previous experiments [83–86].

To model tissue oxygen responses, we simulated a vessel with 3 μm radius and a tissue cylinder of 20 μm radius using Krogh cylinder model.

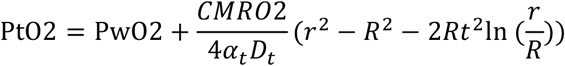

Where, D_t_ = 2800 μm^2^/s [158], a_t_ = 1.39 μM/mmHg [159], CMRO2 = 3 μmole/cm^3^/min [160]. As the transit time of RBCs is much faster than the tissue response time, the observed oxygenation is further smoothed using the response time, which is given by 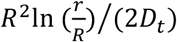 [73]. R is the outer radius and r is the ratio of outer to inner radii. In this way, oxygen delivery from capillaries decays rapidly with distance. As the oxygen probe samples a small region around the tip, we averaged tissue oxygen data within 10μm away from the location of the probe. Finally, to account for the response time of the polarographic oxygen electrodes [20], we smoothed the averaged oxygen trace with a low pass filter.

### White noise, 1/f noise and periodic noise

To test the suitability of power spectral analysis as a method to estimate power-law scales of the same length as the signals used in our analysis, we simulated time series with a stochastic Gaussian process of known long-range temporal dependence (fractional Gaussian noise). The power spectra of different types of signals are shown in **Figure 1**. White noise (**Figure 1**a) has a flat power spectrum whose slope is near 0. Periodic noise (**Figure 1**b) shows a flat spectrum, with the exception of a large bump at 0.2 Hz, the center frequency of the large oscillations. In contrast, 1/f-like noise, generated using a circulant embedding method [161], shows power decreasing with frequency when the power spectrum is plotted on a log-log scale (**Figure 1**c). Summation of a periodic signal with a 1/f-like signal produces a hybrid spectrum (**Figure 1**d).

### Power spectral density and power-law exponent

The power spectrum density (PSD) was obtained using the multitaper technique [103]. As brain activity fluctuations have been observed to have a 1/f power spectrum [162], we tried to fit the power spectrum of oxygen signal with a power-law distribution using least squares fit. However, when linearly-spaced frequency bins are considered under a logarithmic scale, bins in higher-frequencies become progressively denser, and thus gain disproportionate weight with respect to lower frequency-bins in a sub-sequent linear regression. To avoid this potential bias, we upsampled the PSD curve with logarithmically spaced frequency bins, resulting in equally-spaced frequency bins under logarithmic scale, required to properly estimate the spectral exponent. We then used the simple linear regression to the resampled PSD in order to increase comparability to other studies [8, 58].

Although the power spectrum analysis for 1/f-like dynamics estimation is widely used [8, 58, 59, 65, 162], it is important to note that the validity of the applied method has also been criticized [163]. Touboul and Destexhe [163] argue that the power law underlying the 1/f method may not necessarily reflect neurophysiological processes, but could simply arise from purely stochastic mechanisms. Although problematic in testing for the existence of power-law behavior [109], this power spectrum density method gives a good estimate in cases in which the power-law behavior is clear enough [164], or when the aim is at approximating the functional relation rather than setting its exact form, as presented in this study and previous studies [165, 166].

### Control recordings and analyses

Because fluctuations of resistivity in electronic conducting materials also exhibit 1/f noise [108], it is important to demonstrate that our data were not contributed by instrument noise. To address this, we measured PtO_2_ in one dead mouse using the same experimental setup. The power spectrum of these recordings had flat slope (~0) characteristic of white noise (**Figure 2**d).

### Statistical analysis

Statistical analysis was performed using Matlab. All summary data were reported as the mean ± standard deviation (SD) unless stated otherwise. Normality of the samples were tested before statistical testing using Anderson-Darling test (adtest). For comparison of multiple populations, the assumption of equal variance for parametric statistical method was also tested (vartest2 and vartestn). If criteria of normality and equal variance were not met, parametric tests (t test, one-way ANOVA) were replaced with a nonparametric method (Mann-Whiteney U-test, Wilcoxon signed-rank test, Kruskal-Wallis ANOVA). All *p* values were Bonferroni corrected for multiple comparisons. Significance was accepted at *p* < 0.05.

## Supplementary figures

**Supplementary Figure 1.**
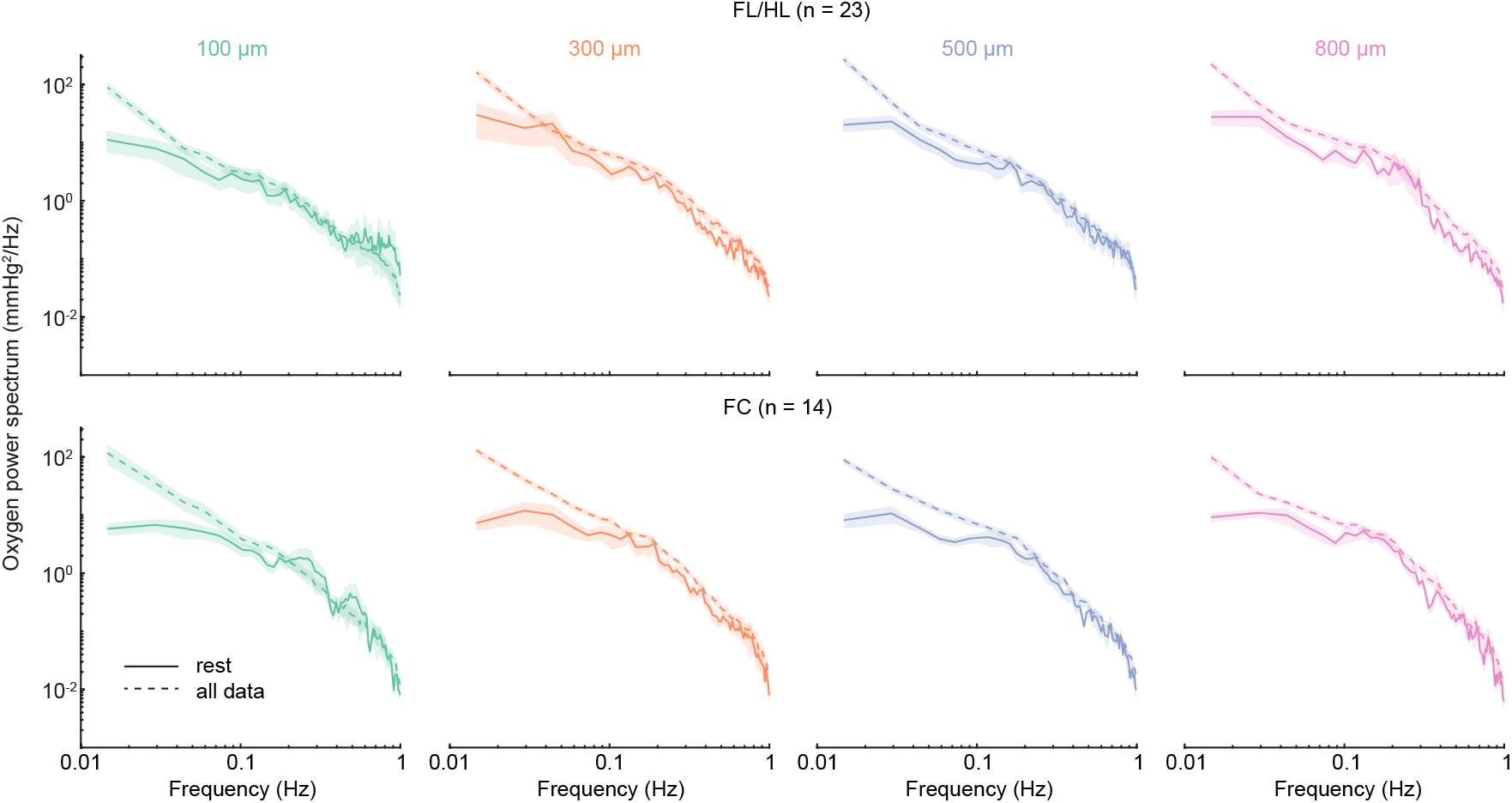
Compare raw spectral power of brain tissue oxygen at rest and including locomotion. Related to **Figure 2**e.

**Supplementary Figure 2.**
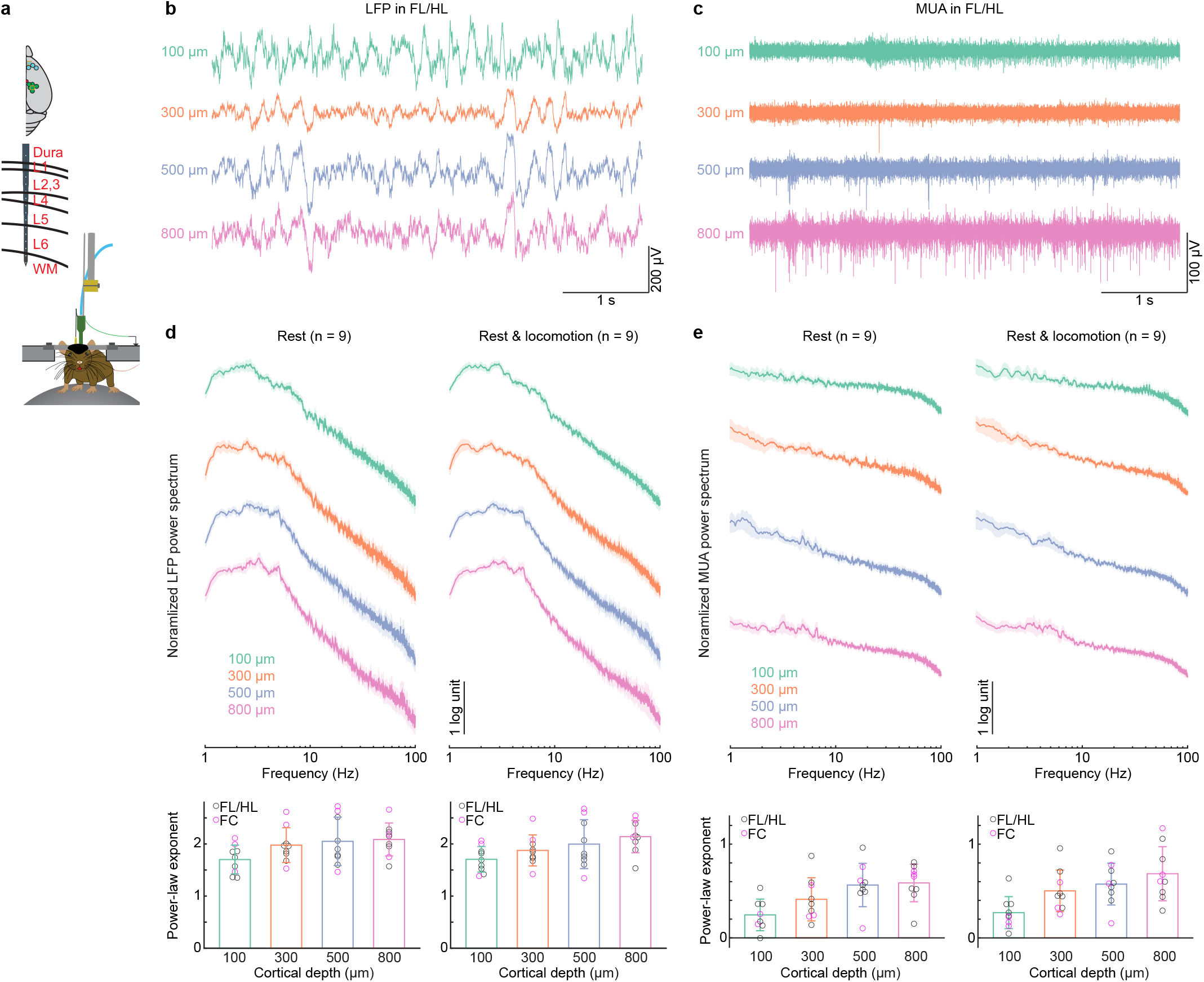
1/f-like power spectra of broadband LFPs and spike rate. (**a**) Experimental setup. Top, schematic showing all laminar electrophysiology measurement sites in FC (n = 3 mice) and FL/HL (n = 6 mice). Bottom, schematic showing the layout of the electrodes and measurement depth. (**b**) An example trace showing the broad-band LFP at different cortical depths in the FL/HL. (**c**) An example trace showing the MUA at different cortical depths in the FL/HL in the same trial as (**b**). (**d**) Top, power spectrum of broadband (1-100 Hz) LFP across cortical depth during rest and periods including rest and locomotion. Bottom, power-law fit exponent. For all cortical layers, the power spectrum showed 1/f-like behavior over 5 Hz, so we fit the power spectrum between 5-100 Hz at each layer with a power-law distribution to avoid the “shoulder”. To compare with the above oxygen measurements, we selected LFPs from recordings 100 μm, 300 μm, 500 μm and 800 μm below the cortical surface. We found the fitted power law exponent was not significantly different across cortical depths at rest (1.70 ± 0.28 at 100 μm, 1.98 ± 0.33 at 300 μm, 2.05 ± 0.46 at 500 μm, 2.09 ± 0.32 at 800 μm; n = 6 mice (9 sites), Kruskal-Wallis test, *χ^2^* (3,35) = 5.89, p = 0.1172). Including locomotion period did not change the relationship across cortical depths (1.71 ± 0.25 at 100 μm, 1.88 ± 0.30 at 300 μm, 2.00 ± 0.47 at 500 μm, 2.14 ± 0.31 at 800 μm; n = 6 mice (9 sites), Kruskal-Wallis test, *χ*^2^(3,35) = 7.55, p = 0.0562). Compared to the resting only data, including the locomotion periods did not increase the power law exponent (average across all cortical layers for each animal, n = 6 mice (9 sites), rest: 1.95 ± 0.37, rest and locomotion: 1.93 ± 0.36, paired t-test, t(35) = 0.9424, p = 0.3525). (**e**) Top, power spectrum of spike train across cortical depths during rest and periods including rest and locomotion. Bottom, power-law fit exponent. In addition to the broad-band LFPs, the spiking rate of neurons is also correlated with vasodilation, increases in blood flow and blood oxygenation [25, 30, 32, 38, 167]. To address whether the spike rate activity also shows 1/f-like activity, we estimated power spectrum of multi-unit spike trains after minimally smoothing the spike trains with a Gaussian kernel with full-width at half maximum (FWHM) of 10 ms. As spikes in surface layers are affected with noise, we made this comparison using the deeper layers, i.e., 500 μm and 800 μm below the pia. Fitting spike train power with a power-law distribution revealed 1/f-like dynamics at rest (power-law exponent: 0.56 ± 0.23 at 500 μm, 0.59 ± 0.20 at 800 μm). The resting power spectrum of spiking activity is much smaller compared to the broad-band LFPs (LFPs: 2.07 ± 0.39, MUA: 0.58 ± 0.21, Wilcoxon rank sum test, p < 0.0001), which is consistent with study [168] showing that 1/f scaling in the LFP power indicates the presence of step like transitions in the LFP trace and says little about properties of the associated neuronal firing. Data are shown as mean ± SEM in (**d**) and (**e**).

**Supplementary Figure 3.**
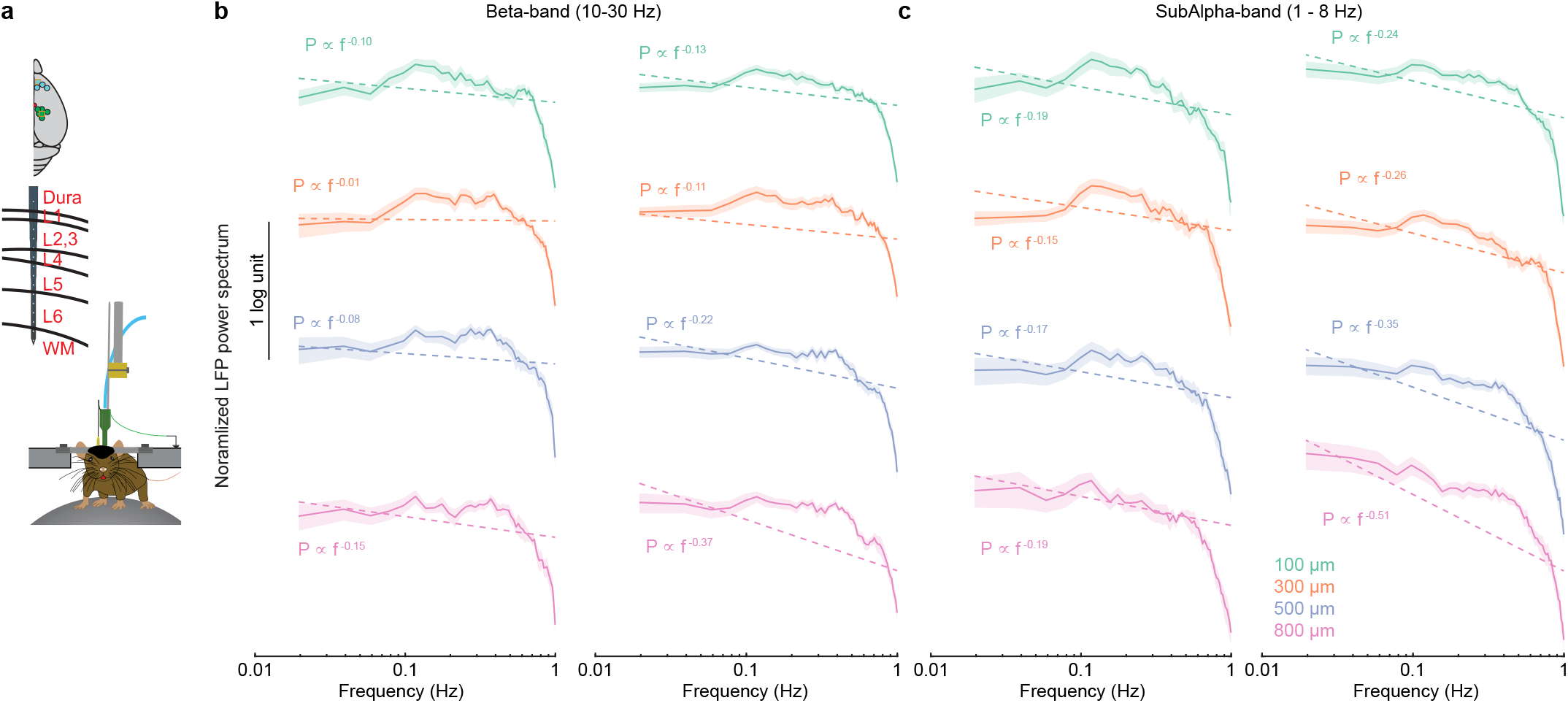
Band-limited power in sub-alpha and beta frequency bands are not 1/f-like. Related to **Figure 3.** (**a**) Experimental setup. (**b**) Power spectrum of the beta-band power of LFP at different cortical depths. Dashed line denotes the linear regression fit of the power. (**c**) As (**b**) but for subalpha-band power of LFP. Data are shown as mean ± SEM.

**Supplementary Figure 4.**
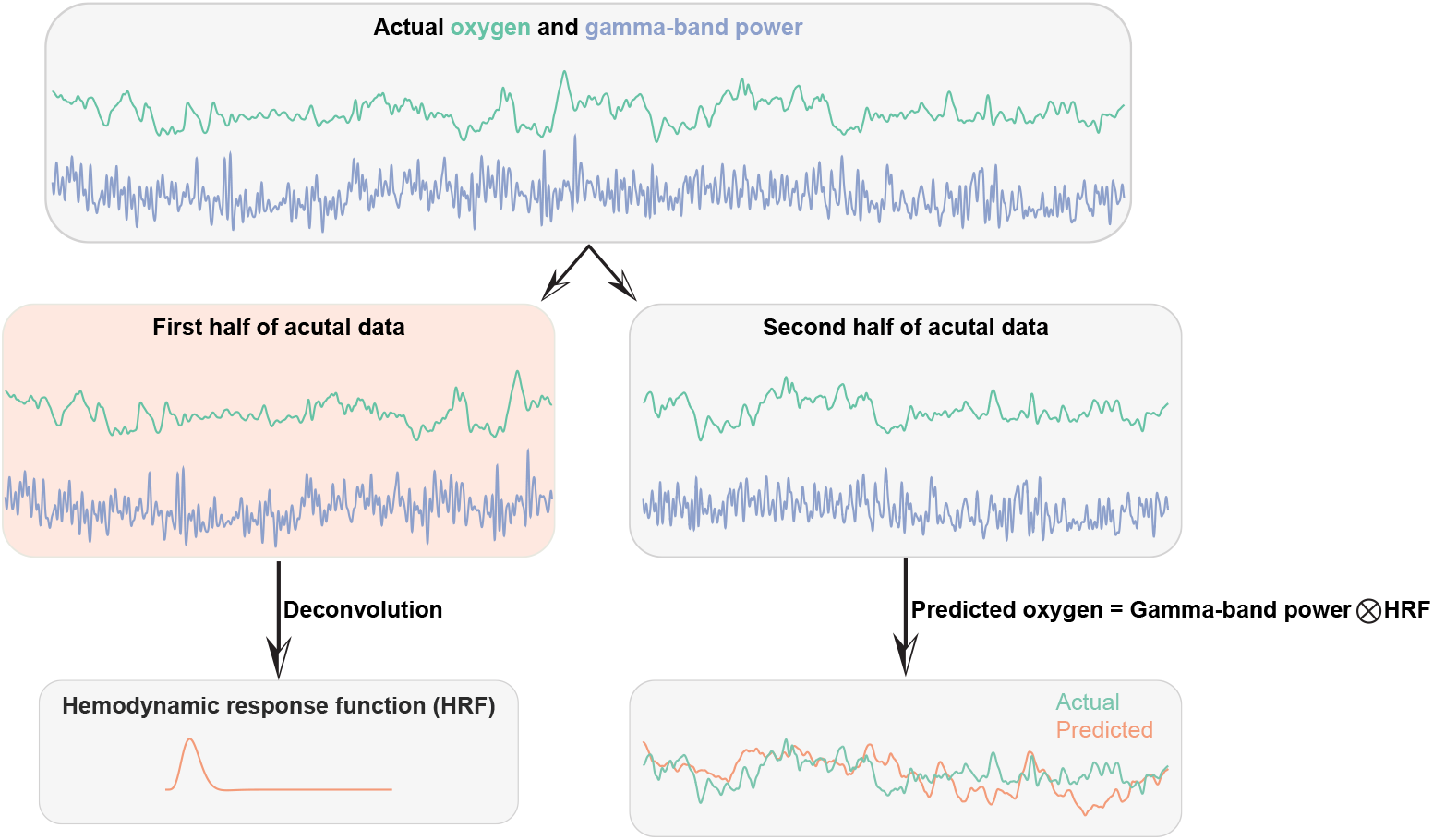
Illustration of the deconvolution and convolution process. Related to **Figure 5.** To estimate how much variance of oxygenation the neural activity can predict, we first split the observed data (~40 min) into two segments with equal length. We then calculated the HRF using the first half of the observed data. To increase the signal-to-noise level of the HRF, we smoothed the HRF using a Savitzky-Golay filter (3^rd^ order, 11-point frame length). Next, we convolved the HRF with the gamma band LFP power from the second half, and estimated the oxygenation predicted by neural activity (orange line).

**Supplementary Figure 5.**
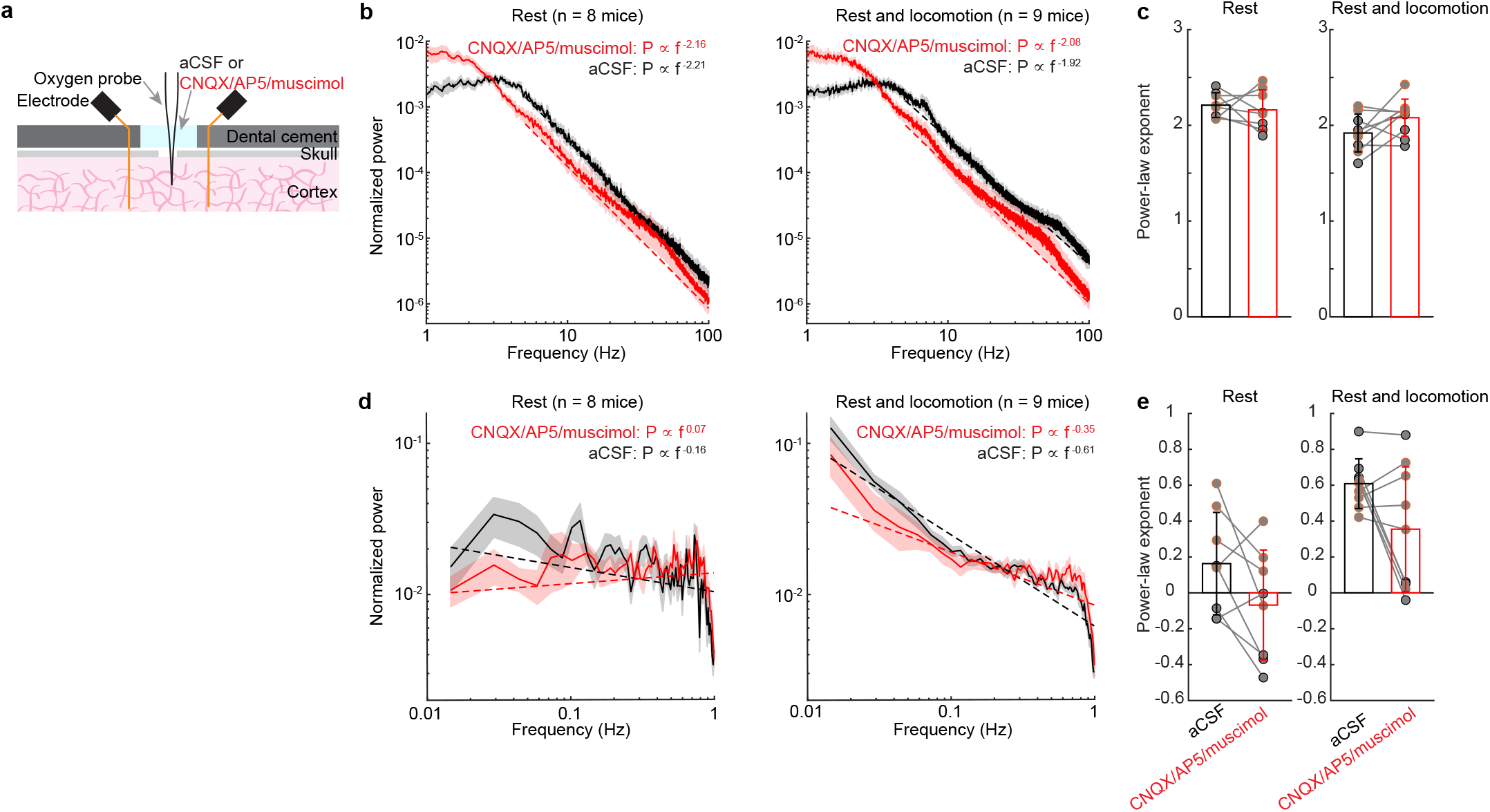
Impact of pharmacological silencing on neural dynamics. Related to **Figure 6.** (**a**) Experimental setup. (**b**) Power spectrum of broadband (1-100 Hz) LFP signal before (black) and after (red) application of CNQX/AP5/muscimol using data during rest (left) and using data including rest and locomotion (right). We asked if this suppression of spiking and LFP oscillations changes the 1/f-like dynamics of neural activity. When the craniotomy was superfused with aCSF, the power spectrum of raw LFP was relatively flat in the lower frequency range (below a “knee” at 5 Hz), and the rest followed a form close to *P* ∝ 1/*f*^2^. Application of CNQX/AP5/muscimol suppressed power at higher frequencies (5-100 Hz), but did not change the power law scaling at rest (aCSF, 2.21 ± 0.13; CNQX/AP5/muscimol, 2.16 ± 0.21; paired t-test, t(7) = 0.5437, p = 0.6035). Including locomotion period did not change the relationship (aCSF, 1.92 ± 0.20; CNQX/AP5/muscimol, 2.08 ± 0.19; paired t-test, t(8) = 1.9318, p = 0.0895). For lower frequency band (1-5 Hz), drug application increased the power law scaling at rest (aCSF, −0.19 ± 0.43; CNQX/AP5/muscimol, 1.40 ± 0.77; paired t-test, t(7) = 9.1504, p < 0.0001). Including running period does not change the relationship (aCSF, −0.15 ± 0.28; CNQX/AP5/muscimol, 1.34 ± 0.69; Wilcoxon rank sum test, p < 0.0001). (**c**) Group average of fitted power-law exponent of broadband LFP. (**d**) Power spectrum of gamma-band power of LFP before (black) and after (red) application of CNQX/AP5/muscimol using data during rest (left) and using data including rest and locomotion (right). We calculated the frequency dependence of the band limited power fluctuations in the gamma-band of LFP during resting periods (aCSF: 0.16 ± 0.29; CNQX/AP5/muscimol: 0.07 ± 0.31, paired t-test, t(7) = 2.0729, p = 0.0769) and during resting and running period (aCSF: 0.61 ± 0.14; CNQX/AP5/muscimol: 0.36 ± 0.35, Wilcoxon rank sum test, p = 0.1903), and did not observe any change in terms of power-law scaling before and after drug application. (**e**) Group average of power-law exponent of BLP for gamma-band.

**Supplementary Figure 6.**
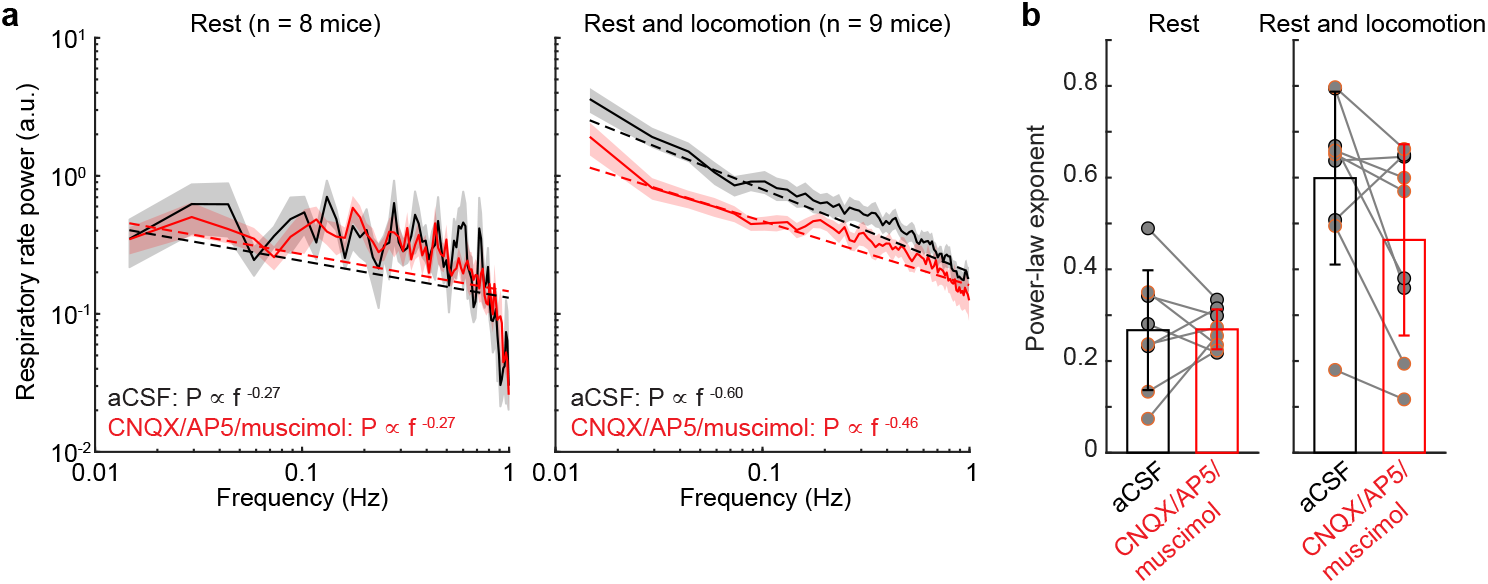
Power spectrum and power law fits of respiratory rate fluctuations. (**a**) Power spectrum of respiratory rate fluctuations before (black) and after (red) application of CNQX/AP5/muscimol using resting data (left) and data including both rest and locomotion (right). (**b**) Power-law exponent of respiratory rate fluctuations before (black) and after (red) application of CNQX/AP5/muscimol using resting data (left) and data including both rest and locomotion (right).

**Supplementary Figure 7.**
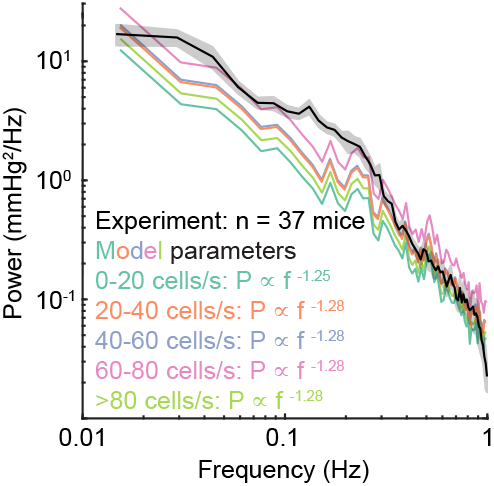
Power spectrum and power law fit of respiratory rate fluctuations. Related to **Figure 7.** Comparison of power spectrum of tissue oxygenation measured using polarographic electrodes (black) and calculated using the simple model under different flow rate. Shaded area denotes 1 SEM.

